# BMP7 functions predominantly as a heterodimer with BMP2 or BMP4 during mammalian embryogenesis

**DOI:** 10.1101/686758

**Authors:** Hyung-Seok Kim, Judith Neugebauer, Autumn McKnite, Anup Tilak, Jan L. Christian

## Abstract

BMP7/BMP2 or BMP7/BMP4 heterodimers are more active than homodimers in vitro, but it is not known whether these heterodimers signal in vivo. To test this, we generated knock in mice carrying a mutation (*Bmp7^R-GFlag^*) that prevents proteolytic activation of the dimerized BMP7 precursor protein. This mutation eliminates the function of BMP7 homodimers and all other BMPs that normally heterodimerize with BMP7. While *Bmp7* null homozygotes are live born, *Bmp7^R-GFlag^* homozygotes are embryonic lethal and have broadly reduced BMP activity. Furthermore, compound heterozygotes carrying the *Bmp7^R-G^* allele together with a null allele of *Bmp2* or *Bmp4* die during embryogenesis with defects in ventral body wall closure and/or the heart. Co-immunoprecipitation assays confirm that endogenous BMP4/7 heterodimers exist. Thus, BMP7 functions predominantly as a heterodimer with BMP2 or BMP4 during mammalian development, which may explain why mutations in either *Bmp4* or *Bmp7* lead to a similar spectrum of congenital defects in humans.

## Introduction

Bone morphogenetic proteins (BMPs) are secreted molecules that were initially discovered as bone inducing factors and were subsequently shown to play numerous critical roles during embryogenesis (Bragdon et al. 2011). Recombinant BMPs are used clinically to treat bone loss caused by trauma or disease, but their usefulness as osteoinductive agents is limited by a short half-life when implanted in vivo (Khan et al. 2012). Understanding how BMP dosage is regulated in vivo is important to prevent congenital birth defects, and to aid in the development of more effective therapeutics to promote bone healing.

BMPs are grouped into subfamilies based on sequence similarity, and can signal as either homodimers or as heterodimers. The class I BMPs, BMP2 and BMP4, can heterodimerize with class II BMPs, consisting of BMPs 5-8 (Guo and Wu 2012). Heterodimers composed of class I and class II BMPs show a higher specific activity than do homodimers. For example, homodimers of BMP2, −4, or-7 can all induce bone formation, but BMP2/7 or BMP4/7 heterodimers are significantly more potent than any homodimer in osteogenic differentiation assays (Aono et al. 1995; Kaito et al. 2018). Likewise, BMP2/6 heterodimers show enhanced ability to activate downstream signaling in embryonic stem cells (Valera et al. 2010). BMP2/7 and BMP4/7 heterodimers also show enhanced ability to induce ventral fate in *Xenopus* and zebrafish (Nishimatsu and Thomsen 1998; Schmid et al. 2000).

While it is widely accepted that recombinant class I/II BMP heterodimers have higher specific activity than homodimers, whether endogenous BMPs function primarily as homodimers or heterodimers in vivo remains controversial. Mutations in either *Bmp2b* or *Bmp7* lead to a complete loss of signaling in zebrafish embryos and this can be rescued by recombinant heterodimers, but not by either homodimer (Little and Mullins 2009). In addition, an ectopically expressed epitope-tagged form of BMP2b pulls down endogenous BMP7 and vice versa (Little and Mullins 2009). Thus, BMP2/7 heterodimers are essential to establish the dorsoventral axis in fish. In *Drosophila*, DPP (the fly homolog of BMP2/4) is not properly localized to the embryonic midline in the absence of SCREW (a BMP7 homolog) and this is proposed to be due to preferential transport of DPP/SCREW heterodimers (Shimmi et al. 2005). However, expression of DPP and SCREW homodimers in distinct regions of the embryo can activate BMP signaling at levels equivalent to the heterodimer (Wang and Ferguson 2005), suggesting that homodimers are sufficient for development.

Evidence that class I/II BMP heterodimers exist or are required for mammalian development is lacking. *Bmp4* or *Bmp2* null homozygotes die during early development with defects in multiple tissues that correlate well with their respective expression domains (Winnier et al. 1995; Zhang and Bradley 1996). Among Class II BMPs, *Bmp8* is restricted to the developing testes and placenta while *Bmp5, Bmp6* and *Bmp7* are broadly expressed throughout embryogenesis (Zhao 2003). Mice homozygous for null mutations in any single Class II *Bmp* gene survive embryogenesis (King et al. 1994; Dudley et al. 1995; Luo et al. 1995; Solloway et al. 1998). By contrast, *Bmp5;Bmp7* or *Bmp6;Bmp7* double mutants are embryonic lethal (Solloway and Robertson 1999; Kim et al. 2001), demonstrating functional redundancy. *Bmp2/7* and *Bmp4/7* double heterozygotes present with no abnormalities or minor skeletal abnormalities (Katagiri et al. 1998), raising the possibility that heterodimers are not required for early mammalian development.

The choice of whether a given BMP will form a homodimer or a heterodimer is made within the biosynthetic pathway. All BMPs are generated as inactive precursor proteins that dimerize and fold within the endoplasmic reticulum (Bragdon et al. 2011). The precursor protein is then cleaved by members of the proprotein convertase (PC) family to generate the active, disulfide bonded ligand along with two prodomain fragments. We have shown that BMP4 and BMP7 preferentially form heterodimers rather than either homodimer when coexpressed in vivo (Neugebauer et al. 2015). *Bmp4 and Bmp7* show overlapping patterns of expression in many tissues (Danesh et al. 2009), suggesting that they may form heterodimers in some contexts. In humans, heterozygous mutations in either *Bmp4* or *Bmp7* are associated with a similar spectrum of ocular, brain and palate abnormalities (Bakrania et al. 2008; Suzuki et al. 2009; Wyatt et al. 2010; Reis et al. 2011), consistent with the possibility that mutations in either gene lead to reduced BMP4/7 heterodimer activity.

BMPs carrying mutations in the PC cleavage motif form inactive dimers with wild type proteins. For example, cleavage mutant forms of BMP7 dominantly interfere with both BMP7 and BMP4 signaling when overexpressed in *Xenopus* (Hawley et al. 1995; Nishimatsu and Thomsen 1998). These studies demonstrate that BMPs can heterodimerize when overexpressed, but do not address whether endogenous BMPs heterodimerize. A mouse carrying a point mutation in the cleavage site of the type II BMP, BMP5, has more severe skeletal abnormalities than *Bmp5* homozygous null mutants (Ho et al. 2008), consistent with the possibility that it interferes with Class I heterodimeric partners, but this has not been explored.

In the current studies, we used genetic and biochemical analysis to test the hypothesis that endogenous heterodimers containing BMP7 exist in vivo. We show that endogenous heterodimers do form in vivo, are the predominant functional ligand in many if not all tissues of developing mouse embryos, and cannot be compensated for by endogenous homodimers. These findings have relevance to understanding the impact of mutations in *Bmp4* or *Bmp7* in humans.

## Results

### *Bmp7^R-GFlag^* homozygotes show earlier lethality and more severe phenotypic defects than *Bmp7* null homozygotes

The phenotypes observed in mice mutant for class I and/or class II *Bmps* can be explained if BMPs function either as homodimers, or as heterodimers (illustrated in Fig. 1A). In the homodimer model, half of the total BMP activity is generated by class I homodimers (BMP2 or 4) and half by the class II homodimers that are broadly expressed in development (BMP5, 6 or 7). In the competing heterodimer model, all of the available class I molecules are covalently dimerized with a class II molecule. A small pool of “excess” class II BMPs form homodimers under wild type conditions, but are available to form heterodimers to compensate for loss of any single class II BMP. These predictions are consistent with our findings that BMPs preferentially form heterodimers (Neugebauer et al. 2015), and with genetic data showing greater redundancy among class II BMPs than among class I BMPs.

**Figure 1.**
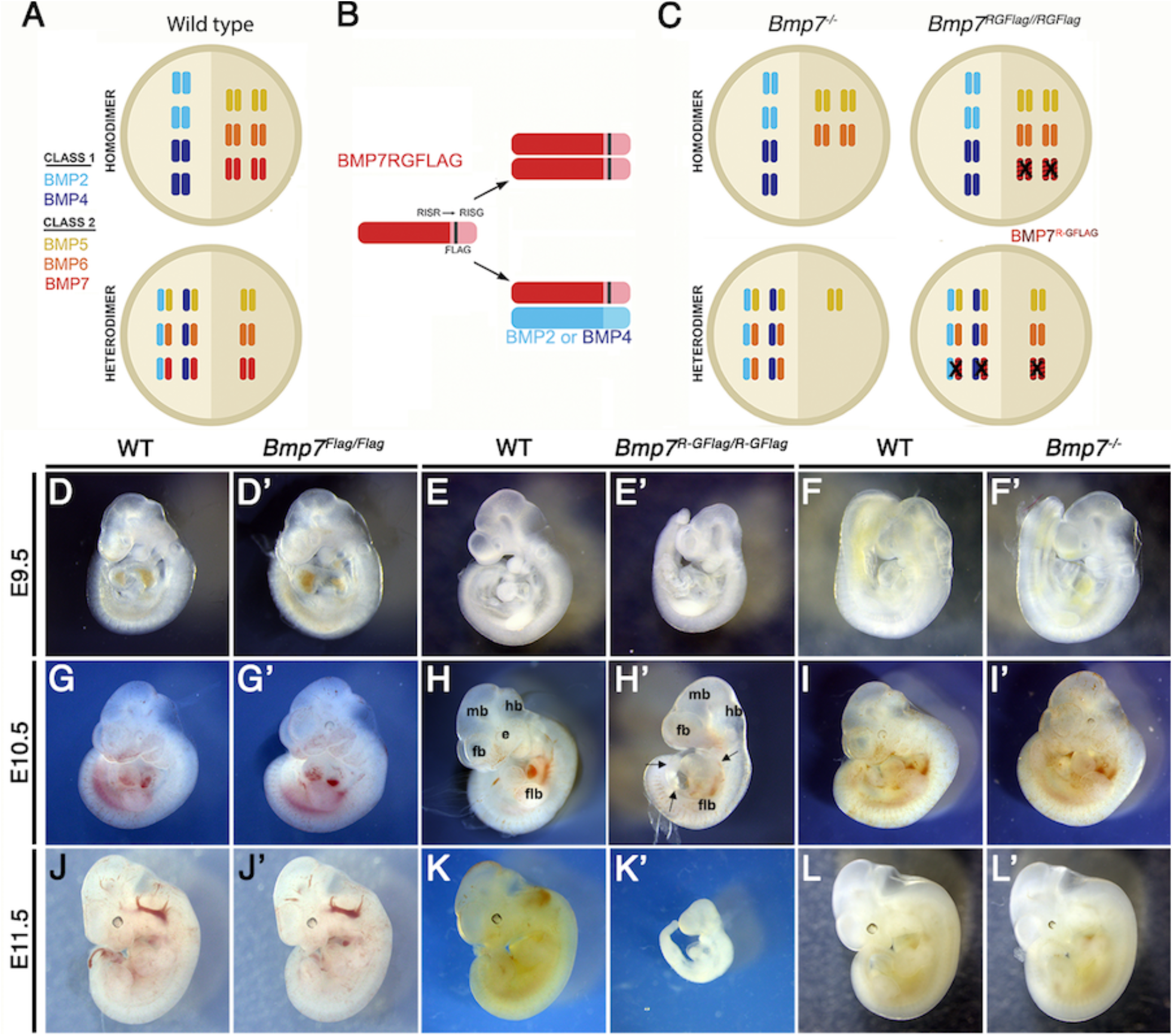
*Bmp7^R-GFlag^* homozygotes show earlier lethality and more severe phenotypic defects than *Bmp7* null homozygotes. (A) Illustration of total BMP activity in a hypothetical tissue if class I and II BMPs function predominantly as homodimers (top) or heterodimers (bottom). Model assumes class I and class II BMPs each contribute 50% of total activity, but there are more class II ligands, and thus greater redundancy. In the heterodimer model, it is assumed that class I/II heterodimers form preferentially and that there is an excess of class II BMPs that form homodimers. (B) Illustration of BMP7R-GFlag precursor protein forming homodimers (top) or heterodimers with BMP2 or BMP4 (bottom). Prodomain: dark shading, mature domain: light shading, black bar: FLAG epitope. (C) Illustration showing the predicted fraction of BMP activity that will be lost in *Bmp7^R-GFlag^* or *Bmp7* null homozygotes if BMPs function predominantly as homodimers (top) or as heterodimers (bottom). In the homodimer model, there is an equivalent reduction in BMP activity in *Bmp7^−/−^* or *Bmp7^R-GFlag/Flag^* mice because only BMP7 homodimers are absent or inactive (black X), respectively. In the heterodimer model, excess class II molecules that normally form homodimers will fill in to maintain the heterodimer pool in *Bmp7* null mutants, but cannot do so in *Bmp7^R-GFlag^* mutants. This leads to a greater reduction in the more active heterodimer pool and in total BMP activity. (D-L’) Photograph of E9.5-11.5 (age indicated to left of each row) wild type (D-L) or mutant (D’-L’; genotype listed at top of each column) littermates. A minimum of eight embryos of each genotype were analyzed at each stage. Arrows in H’ indicate pericardial edema. fb; forebrain, mb; midbrain, hb; hindbrain, flb; forelimb bud, e; eye.

To ask whether heterodimers containing endogenous BMP7 are required for normal development, we generated a *Bmp7* cleavage mutant mouse (*Bmp7^R-GFlag^*) (Supplemental Fig. S1). These mice have a point mutation that changes the amino acid sequence of the PC cleavage motif from -RISR- to -RISG-, as well as sequence encoding a Flag epitope tag knocked in to the *Bmp7* allele (Fig. 1B). *Bmp7^R-GFlag^* mice express endogenous levels of a non-cleavable, inactive BMP7 precursor protein. A control mouse that carries the Flag-epitope tag at the wild type *Bmp7* locus (*Bmp7^Flag^*) was generated in parallel.

If BMP7 signals exclusively as a homodimer, then *Bmp7^R-GFlag^* homozygotes should show the same reduction in BMP activity as *Bmp7* null mutants (illustrated in Fig. 1B-C; upper diagrams), and would be predicted to die shortly after birth due to kidney defects (Dudley et al. 1995; Luo et al. 1995). By contrast, if class I/II heterodimers are the primary functional ligand, then the BMP7R-GFlag precursor protein would form non-functional covalent heterodimers with all endogenous class I BMPs with which it normally interacts (Fig. 1B, lower diagram). In this case, the “surplus pool” of class II BMPs that can buffer the heterodimer pool in *Bmp7* null mutants (Fig. 1C, lower left) will be unable to compensate in *Bmp7^R-GFlag^* mutants (Fig. 1C, lower right). This would lead to a greater reduction in the heterodimer pool, lower total BMP activity and more severe phenotypic defects in *Bmp7^R-GFlag^* than in then *Bmp7* null mutants.

*Bmp7^R-GFlag/+^*, *Bmp7^Flag/+^* or *Bmp7^-/+^* mice were intercrossed to determine viability. *Bmp7^Flag/Flag^* embryos were recovered at the predicted Mendelian frequency throughout development and were adult viable with no apparent defects (Table 1A). *Bmp7* null homozygotes were recovered at the predicted Mendelian ratio between embryonic day (E)9.5 and E18.5 (Table 1B) but died shortly after birth. By contrast, *Bmp7^R-GFlag/R-GFlag^* mice were present at the predicted Mendelian frequency through E11.5 but were not recovered after this stage (Table 1C).

**Table 1A.** Progeny from *Bmp7^Flag/+^* intercrosses

| Age | *Bmp7*<sup>+/+</sup> | *Bmp7*<sup>Flag/+</sup> | *Bmp7*<sup>Flag/Flag</sup> | Total |
| --- | --- | --- | --- | --- |
| P28 | 8 (20%) | 24 (60%) | 8 (20%) | 40 |
| E9.5-14.5 | 23 (31%) | 34 (46%) | 17 (23%) | 74 |

Data are presented as number (percent).

**Table 1B.**
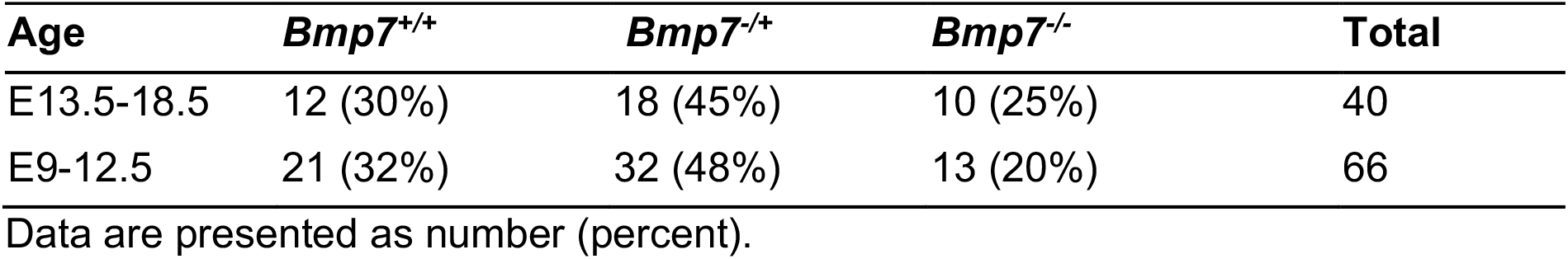
Progeny from *Bmp7^-/+^* intercrosses

**Table 1C.**
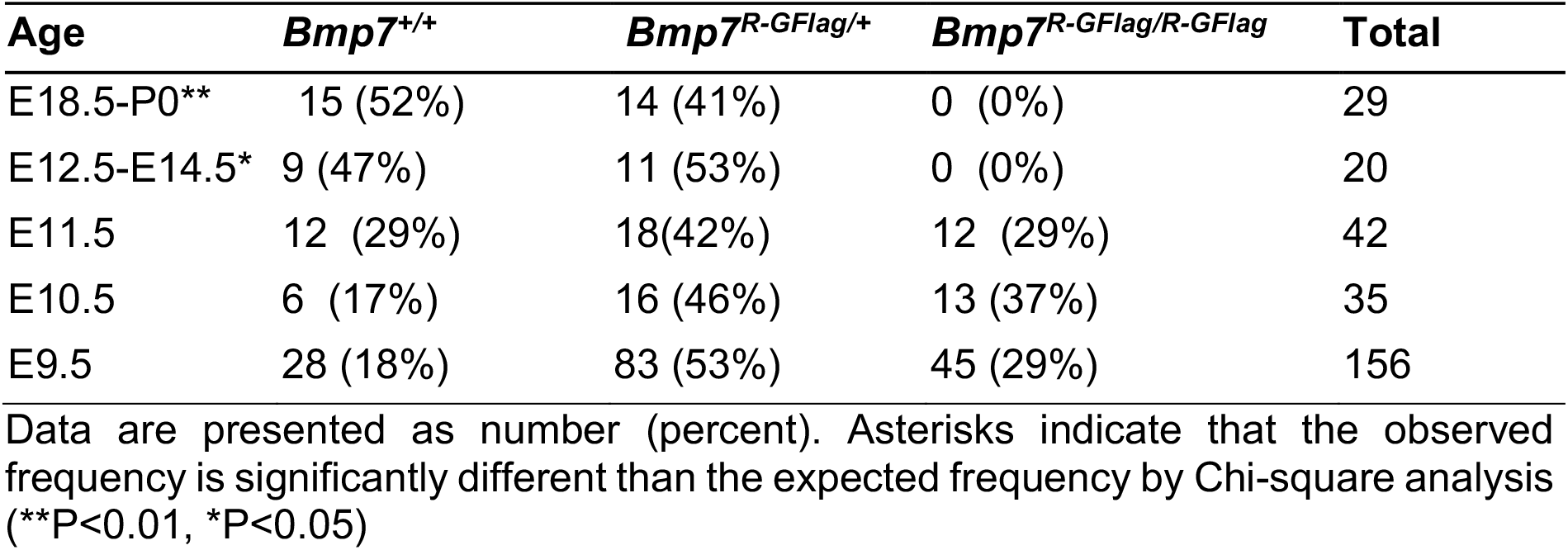
Progeny from *Bmp7^R-GFlag^+* intercrosses

| Age | *Bmp7*<sup>+/+</sup> | *Bmp7*<sup>R-GFlag/+</sup> | *Bmp7*<sup>R-GFlag/R-GFlag</sup> | Total |
| --- | --- | --- | --- | --- |
| E18.5-P0** | 15 (52%) | 14 (41%) | 0 (0%) | 29 |
| E12.5-E14.5* | 9 (47%) | 11 (53%) | 0 (0%) | 20 |
| E11.5 | 12 (29%) | 18(42%) | 12 (29%) | 42 |
| E10.5 | 6 (17%) | 16 (46%) | 13 (37%) | 35 |
| E9.5 | 28 (18%) | 83 (53%) | 45 (29%) | 156 |

Data are presented as number (percent). Asterisks indicate that the observed frequency is significantly different than the expected frequency by Chi-square analysis (\*\*P<0.01, \*P<0.05)

*Bmp7^Flag/Flag^* (Fig. 1D, D’, G, G’, J, J’) and *Bmp7*^−/−^ (Fig. 1F, F’, I, I’, L, L’) embryos were indistinguishable from wild type littermates at E9.5-11.5, with the exception of slightly smaller eyes in 25% of the *Bmp7*^−/−^ embryos examined at E11.5 (*n*=7; Fig. 1L, L’). *Bmp7^R-GFlag/ R-GFlag^* embryos appeared grossly normal but were slightly smaller, and had smaller limb buds than age matched (by somite number) wild type littermates at E9.5 (*n*=45; Fig. 1E, E’, Supplemental Fig. S2). *Bmp7^R-GFlag^* homozygotes were smaller than littermates at E10.5 (*n*=13; Fig. 1H, H’) and were resorbing at E11.5 (*n*=12; Fig. 1K, K’). All *Bmp7^R-GFlag^* homozygotes showed multiple abnormalities at E10.5 (Fig. 1H, H’) including smaller and less distinct forebrain (fb), midbrain (mb) and hindbrain (hb), pericardial edema (arrows), smaller limb buds (flb) and no eye (e). Thus, expression of wild type levels of an uncleavable BMP7 precursor protein leads to earlier lethality and more severe phenotypic defects than does complete absence of BMP7 protein, suggesting that endogenous BMP7-containing heterodimers perform essential functions during early embryogenesis.

*Bmp7^R-GFlag^* heterozygotes were adult viable (Table S1), but 23% showed runting, microphthalmia and/or anophthalmia as early as E14.5 (*n*=13, Supplemental Fig. S3A-B) that persisted into adulthood (Supplemental Fig S3, E-F). These defects were never observed in *Bmp7* null heterozygotes (*n*=8; Supplemental Fig. S3C) although one or both eyes were absent in late gestation *Bmp7* null homozygotes (*n*=5; Fig. S3D) as previously reported (Dudley et al. 1995; Luo et al. 1995). Skeletal analysis revealed that the fibula was shortened and failed to articulate with the knee in 13% of *Bmp7^R-GFlag^* heterozygotes that were analyzed (*n*=14; Supplemental Fig. S2G, H). This defect has been observed in mice in which both *Bmp2* and *Bmp7* are conditionally deleted from the limbs, but not in mice lacking any single BMP family member (Bandyopadhyay et al. 2006). These findings support a model in which the BMP7R-GFlag precursor protein dominantly sequesters endogenous class I BMPs in non-functional dimers.

### *Bmp7^RGFIaa^* homozygotes show defects in heart development that are absent in *Bmp7* null homozygotes

*Bmp2, 4, and 7* are expressed in overlapping domains of the developing heart (Dudley and Robertson 1997; Danesh et al. 2009), raising the possibility that heart defects cause embryonic lethality in *Bmp7^R-GFlag/R-GFlag^* mutants. At E10.5, hearts dissected from *Bmp7^Flag^* (*n*= 5) and *Bmp7* null homozygotes (*n*= 6) were indistinguishable from wild type littermates with morphologically distinguishable atria, ventricles, and outflow tract (OFT) (Fig. 2A-A’, C-C’). By contrast, all hearts of *Bmp7^R-GFlag/R-GFlag^* embryos (*n*=6) appeared to have thinner walls, had a single common atrium, a smaller right ventricle and a small, malformed OFT relative to wild type littermates (Fig. 2B-B’). In addition, although hearts of *Bmp7^R-GFlag/R-GFlag^* mutants were morphologically normal at E9.5, expression of the direct BMP target gene, *Nkx2.5* (Lien et al. 2002), was severely reduced in all hearts of *Bmp7^R-GFlag/R-GFlag^* mutants relative to littermates at E9.5 (*n*=10) and E10.5 (*n*=6) (Fig. 2E-E’, H-H’, K-K’, N-N’). No differences were detected in expression of *Nkx2.5* in *Bmp7^Flag^* (Fig. 2D-D’, G-G’, J-J’, M-M’) or in *Bmp7* null homozygotes (Fig. 2F-F’, I-I’, L-L’, O-O’) relative to wild type littermates.

**Figure 2.**
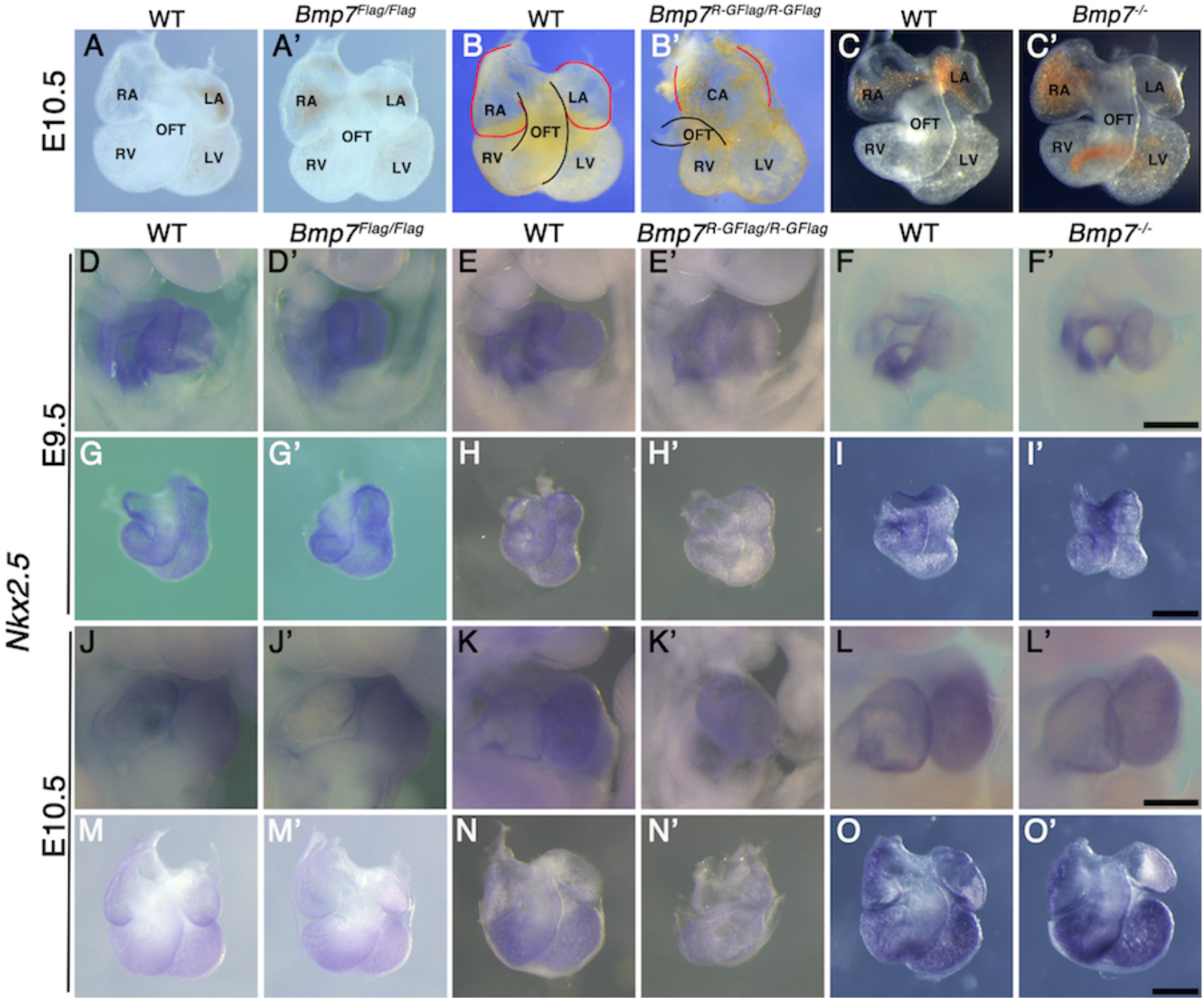
*Bmp7^R-GF!ag^* homozygotes show defects in heart development that are absent in *Bmp7* null homozygotes. (A-C’) Photographs of hearts dissected from E10.5 mice carrying targeted alleles of *Bmp7* (A’-C’) and wild type littermates (A-C). Genotypes are indicated above each panel. The OFT is outlined in black, and the atrium in red in B and B’. (D-O’) Expression of *Nkx2.5* was analyzed by whole mount in situ hybridization in mice carrying targeted alleles of *Bmp7* (D’-O’) and wild type littermates (D-O) at E9.5 or E10.5 as indicated to the left of each row. Genotypes are indicated above each panel. Close up photographs of hearts in intact embryos (D-F’, J-L’) or dissected free of embryos (G-I’, M-O’) are shown. Scale bars in all panels correspond to 500 μM.

### *Bmp7^R GFIaa^* mutants, but not *Bmp7* null mutants, show reduced BMP activity in multiple tissues

To test whether BMP activity is more severely reduced in *Bmp7^R-GFlag^* mutants than it is in *Bmp7* null mutants, we analyzed BMP activity in *BRE:LacZ* transgenic embryos at E9.5, before gross morphological abnormalities are detected in *Bmp7^R-GFlag^* homozygotes. This transgene contains a BMP-responsive element coupled to LacZ, which serves as an in vivo reporter of BMP signaling downstream of all endogenous BMP ligands (Monteiro et al. 2004). X-GAL staining for ß-galactosidase activity in *Bmp7^+/+^;BRE:LacZ* embryos revealed strong endogenous BMP activity in the brain, eye, branchial arches (BA), heart, and ventroposterior mesoderm (VPM) (Fig. 3A, C). No differences in BMP activity were detected in any of these tissues in *Bmp7*^−/−^ embryos (Fig. 3A, B). By contrast, as shown in Fig. 3C and D, *Bmp7^R-GFlag/R-GFlag^* embryos exhibited a reproducible reduction in BMP activity in the brain, ventroposterior mesoderm, and heart. The reduction in staining in the heart of *Bmp7^R-GFlag/R-GFlag^* embryos was most pronounced in the right ventricle (outlined in white in the inset) and the OFT (outlined in yellow in inset). In addition, staining was completely absent in the eye (Fig. 3D). Thus, mice expressing endogenous levels of an uncleavable form of the BMP7 precursor protein show widespread loss of BMP activity that is not observed in mice lacking BMP7 protein.

**Figure 3.**
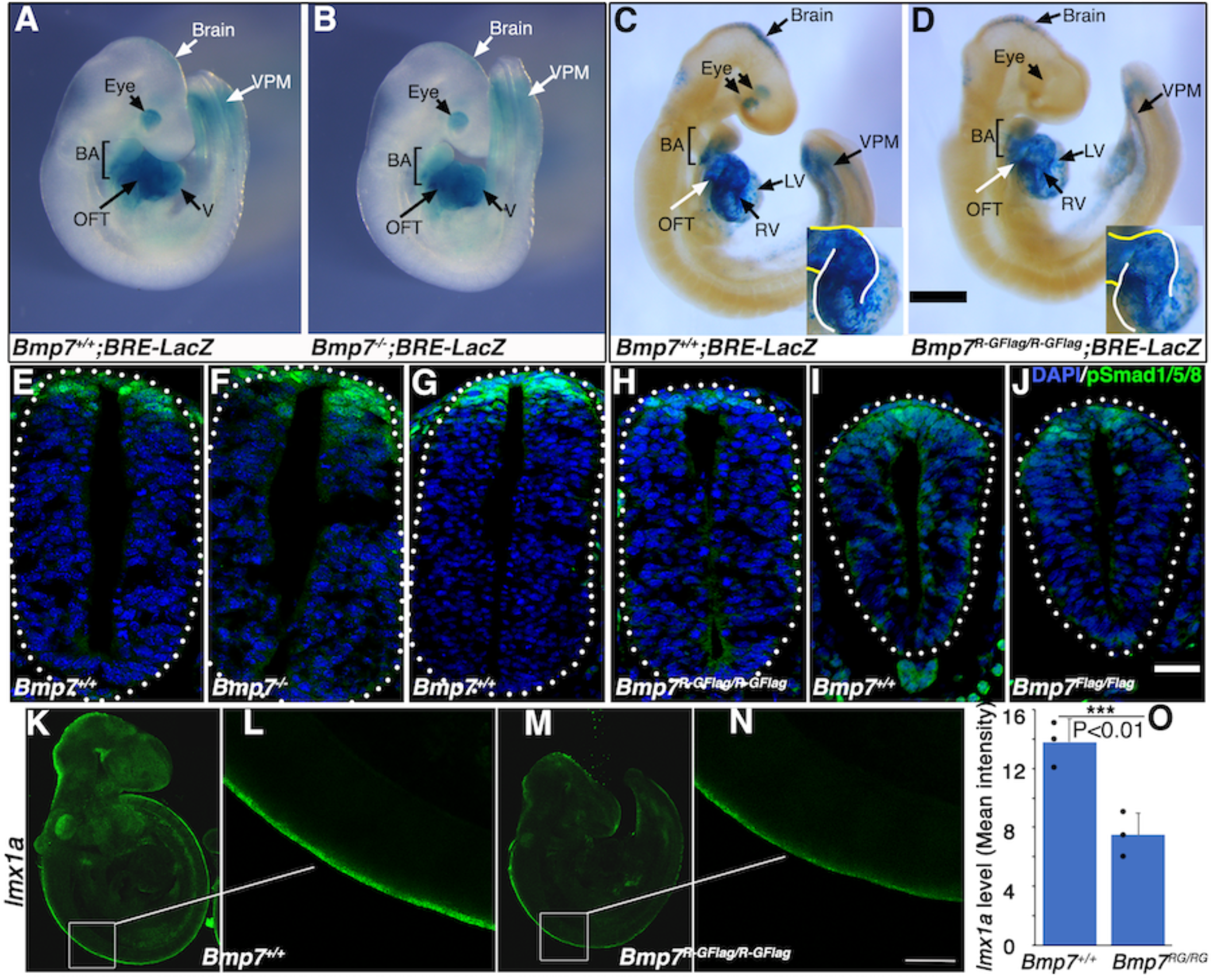
*Bmp7^R-GFlag^*mutants, but not *Bmp7* null mutants, show reduced BMP activity in multiple tissues. (A-D) E9.5 *Bmp7*^−/−^ (B), *Bmp7^R-GFlag/R-GFlag^* (D) and wild type littermates (A, C) carrying a BRE-LacZ transgene were stained for ß-galactosidase activity to detect endogenous BMP pathway activation. Embryos from a single litter were stained for an identical period of time under identical conditions. A minimum of three embryos of each genotype were examined. VPM; ventral posterior mesoderm, BA; branchial arches, OFT; outflow tract, RV; right ventricle, LV; left ventricle. Insets in C, D show an enlarged view of the heart with the OFT and RV outlined in yellow and white, respectively. In C, left and right eyes are visible in the cleared embryo. Scale bars correspond to 1 mm. (E-H) Transverse sections through the spinal cord of E9.5 *Bmp7*^−/−^(F), *Bmp7^R-GFlag/R-GFlag^* (H), *Bmp7^Flag/Flag^* (J), and wild type littermates (E, G, I) were immunostained with antibodies specific for pSmad1/5/8 and nuclei were stained with DAPI. Dorsal is up. Scale bars correspond to 20 μm. Three embryos of each genotype were analyzed. (K-N) *In situ* HCR was used to analyze expression of *Imx1a* in E9.5 *Bmp7^R-GFlag/R-GFlag^* (M-N) and wild type littermates (K-L). White boxes in K and M indicate the region of staining shown in L and N. Scale bar corresponds to 100 μm. (O) Levels of *Imxa1* transcripts (mean fluorescence ± s.d., data analyzed by two tailed t-test). Fluorescence intensity was quantity in comparable regions of the spinal cord in three embryos of each genotype, dots represent image intensity for each embryo that was scored.

*Bmp2, 4* and *7* are co-expressed in the dorsal surface ectoderm overlying the spinal cord by E8 (Solloway and Robertson 1999; Danesh et al. 2009). BMP signaling from the ectoderm is required for induction of the roof plate at E9, and BMPs and other factors secreted from the roof plate are subsequently required for specification, migration and axon guidance of dorsal interneurons (Chizhikov and Millen 2005). To analyze BMP activity in this important signaling center, we immunostained sections of E9.5 embryos using an antibody specific for the active, phosphorylated form of BMP pathway-specific SMADs (pSmad1/5/8). Levels of pSmad1/5/8 were unchanged in the roof plate of *Bmp7^−/−^* (Fig. 3E-F) and *Bmp7^Flag/Flag^* embryos relative to wild type littermates (Fig. 3I-J), but were severely reduced in the roof plate of *Bmp7^R-GFlag^* homozygotes (Fig. 3G-H).

To further test whether BMP heterodimers are required for initial induction of the roof plate, we analyzed expression of *lmx1a* using *in situ* hybridization chain reaction (HCR). Expression of *lmx1a* is induced in the nascent roof plate downstream of BMP signaling from the epidermal ectoderm, and is the major mediator of BMP signaling in the dorsal neural tube (Chizhikov and Millen 2004). Expression of *lmx1a* was reduced by 50% in all E9.5 embryos that were examined (*n*=3; Fig. 3K-O). Thus, BMP7 containing heterodimers secreted from the epidermal ectoderm are major contributors to roof plate induction.

### Bmp7R-GFlag homodimers do not act outside of cells to block BMP activity

Our results support a model in which BMP7-containing class I/II heterodimers are the predominant functional ligand in early embryos. An assumption of this model is that the BMP7R-GFlag precursor protein forms covalent heterodimers with class I BMP precursor proteins inside of cells in which they are co-expressed (illustrated in Fig. 1B), thus sequestering heterodimers in non-functional complexes that are unable to activate their receptors. An alternate possibility is that BMP7R-GFlag precursor forms uncleavable homodimers that are secreted and form non-functional complexes with BMP receptors on the cell surface, thereby blocking the ability of class I BMP homodimers to activate their cognate receptors. To test this possibility, we expressed BMP7R-GFlag or BMP4 in HEK293 cells, and collected BMP7R-GFlag precursor protein or mature BMP4 that was secreted into the culture medium. Non-transfected HEK293 cells were then exposed to equivalent amounts of mature BMP4 alone, BMP7R-GFlag precursor alone, or both together for one hour prior to analyzing levels of pSmad1/5/8 by immunoblot (illustrated in Fig. 4A). Cells incubated with BMP7R-GFlag showed the same barely detectable level of immunoreactive pSmad1/5/8 as did control cells (Fig. 4B, compare lane 1 and 3), indicating that the precursor protein lacks activity. Levels of pSmad1/5/8 were elevated to the same extent in cells incubated with BMP4 alone, or with BMP4 and BMP7R-GFlag together (compare lane 2 and 4). Thus, uncleaved BMP7R-GFlag precursor protein homodimers cannot act outside of the cell to block the ability of BMP4 to signal.

**Figure 4.**
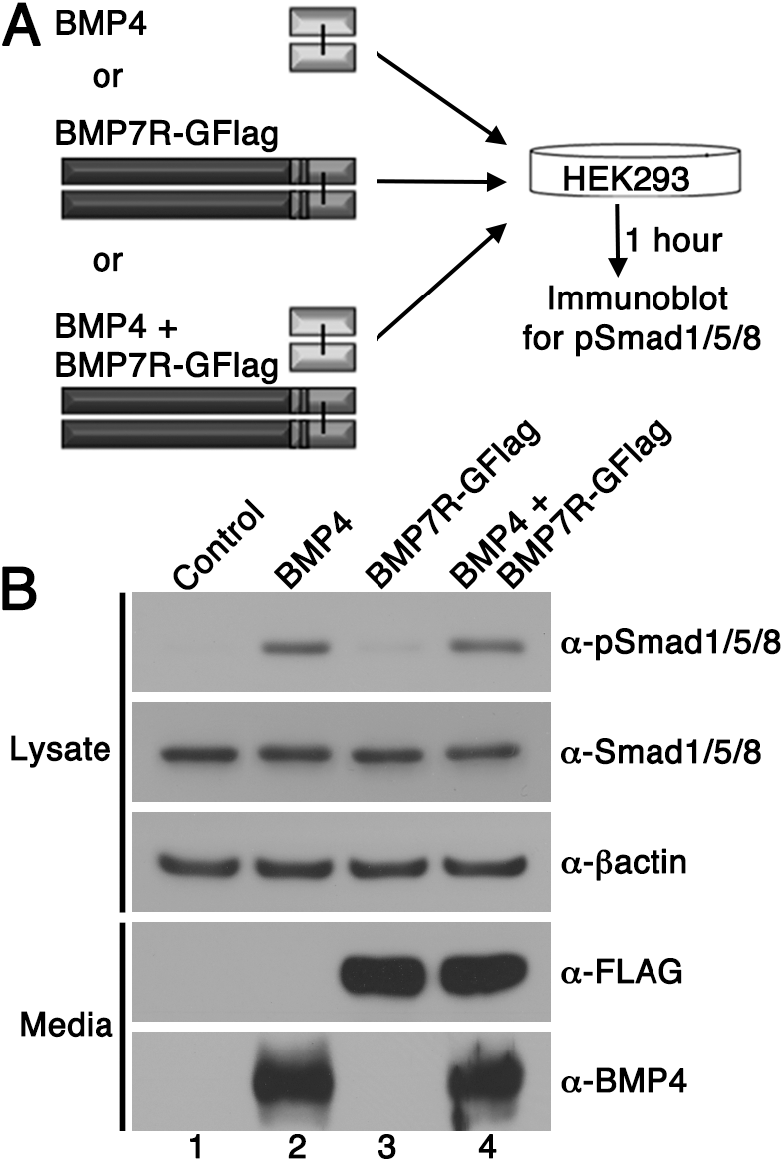
BMP7R-GFlag homodimers do not act outside of cells to block BMP signaling. HEK293 cells were cultured for one hour in conditioned media containing equivalent amounts of BMP4 ligand alone, BMP7R-GFlag precursor protein alone or BMP4 ligand and BMP7R-GFlag precursor together as illustrated. Levels of pSmad1/5/8, total Smad1/5/8, BMP7R-GFlag and BMP4 were analyzed by immunoblot. Blots were reprobed for ß-actin as a loading control. Results were reproduced in three independent experiments.

### Analysis of compound heterozygotes demonstrates that BMP2/7 and BMP4/7 heterodimers are functionally important during early development

To further test the idea that heterodimers of BMP7 together with BMP2 and/or BMP4 are essential for embryogenesis, and to ask which class I ligand(s) contribute to distinct developmental processes, we generated compound heterozygous mutants that carry one copy of the *Bmp7^R-GFlag^* allele in combination with a null allele of *Bmp2* or *Bmp4*. The heterodimer model predicts that removing a single copy of *Bmp2* or *Bmp4* will reduce the heterodimer pool, leading to a modest reduction in total BMP activity (illustrated at top of Fig. 5). A further prediction is that the additional removal of a single copy of *Bmp7* (*Bmp2^-/+^;Bmp7^-/+^* or *Bmp4^-/+^;Bmp7^-/+^* compound mutants) will not lead to further depletion of the heterodimer pool, due to the ability of other class II BMPs to substitute for BMP7. Consistent with this prediction, *Bmp2, 4 or 7* null heterozygotes are adult viable and show mild skeletal defects that are not substantially worse in *Bmp2^-/+^;Bmp7^-/+^* or Bmp4^-/+^;Bmp7^-/+^compound heterozygotes (Katagiri et al. 1998). The heterodimer model predicts a different outcome in the case of *Bmp2^-/^+;Bmp7^R-GFlag/+^* or *Bmp4^-/+^;Bmp7^R-GFlag/+^* mice, since the BMP7R-GFlag protein sequesters a fraction of the available BMP2 and/or BMP4 in non-functional heterodimers (model, far right) such that a further reduction in *Bmp2* or *Bmp4* gene dosage will cause additional loss of the heterodimer pool (model, far right).

**Figure 5.**
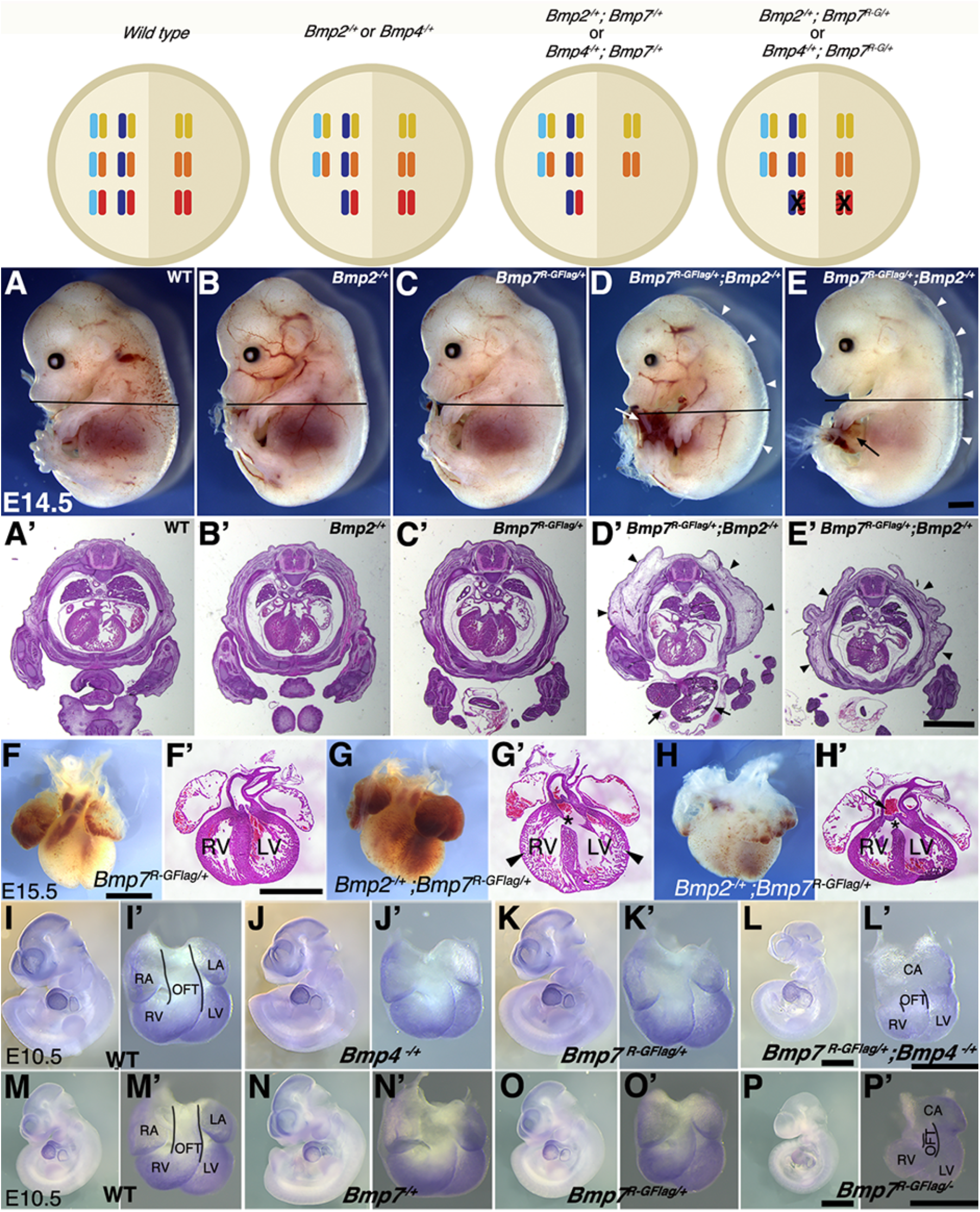
Analysis of compound heterozygotes shows that BMP2/7 and BMP4/7 heterodimers are required for early development. Schematic illustration of total BMP activity (represented by circle) in a hypothetical tissue in embryos carrying wild type or mutant alleles of *Bmp2, 4* and/or 7. Model assumes all available BMP2 and BMP4 molecules are heterodimerized with a class II ligand and there is an excess pool of homodimeric class II ligands. In embryos lacking a single copy of *Bmp2* or *Bmp4*, activity contributed by the heterodimer pool is reduced but there is no further reduction in heterodimers when a single copy of *Bmp7* is also removed, due to redundancy with other Class II BMPs. A fraction of the heterodimer pool is inactivated in *Bmp7^R-GFlag^* heterozygotes and additional removal of a single copy of *Bmp2* or *Bmp4* further depletes the heterodimer pool. (A-E’) Photographs of E14.5 wild type and mutant littermates (A-E) and corresponding hematoxylin and eosin stained transverse sections through the abdomen (A’-E’; approximate position of section indicated by the black bar in A-E). Arrows indicated externalized viscera (D, D’, E) and arrowheads denote peripheral edema (D, D’, E, E’). (F-H’) Ventral views of hearts dissected from E15.5 *Bmp7^R-GFlag/+^* (F) or *Bmp7^R-GFlag/+^;Bmp2^-/+^* embryos (G-H) and corresponding hematoxylin and eosin stained coronal sections (F’-H’). Asterisks denote VSDs (G’, H’), arrow indicates abnormal positioning of the aorta exiting the right ventricle (H’), arrowheads indicate thin, non-compacted ventricular wall (G’). (I-P’) Expression of *Nkx2.5* was analyzed by whole mount in situ hybridization in littermates generated by intercrossing *Bmp4^-/+^* and *Bmp7^R-GFlag/+^* (I-L’) or *Bmp7^-/+^* and *Bmp7^R-GFlag/+^* mice (M-P’). Photographs of intact embryos at E10.5 (I-P) and photographs of hearts dissected from corresponding embryos (I’-P’) are shown. RA; right atrium, LA; left atrium, CA; common atrium, RV; right ventricle, LV; left ventricle, OFT; outflow tract. The OFT is outlined in I’, L’, M’ and P’. Scale bars in all panels correspond to 1 mm.

When *Bmp7^R-GFlag/+^* and *Bmp2^/+^* mice were intercrossed, *Bmp2^-/^+;Bmp7^R-GFlag/+^* mutants were present at the predicted Mendelian ratios at E15.5, but were not recovered at weaning (Table 2A). *Bmp2^-/+^* and *Bmp7^R-GFlag/+^* mice were indistinguishable from wild type littermates (Fig. 5A-C), whereas all (*n*=10) E14.5 *Bmp2^/+^;Bmp7^R-GFlag^* embryos showed peripheral edema (Fig. 5D-E, D’-E’, arrowheads) along with defects in ventral body wall closure that ranged from umbilical hernia (Fig. 5 E, arrow, *n*=4) to omphalocele, in which the liver and other visceral organs were externalized (Fig. 5D,D’, arrows, *n*=6). In addition, six of the ten compound heterozygotes were runted (Fig. 5 D). Peripheral edema is often associated with cardiovascular defects and thus we examined the hearts of E15.5 embryos. The heart from one *Bmp2^/+^;Bmp7^R-GFlag/+^* embryo was much smaller than that of littermates and appeared atrophied (Supplemental Fig. S4A,B). Out of four *Bmp2^/+^;Bmp7^R-GFlag/+^* hearts that were examined histologically, three showed ventricular septal defects (VSDs) (Fig. 5G’,H’, asterisks), and two showed defects in alignment of the aorta and pulmonary trunk (Fig. 5G’,H’) relative to wild type or single mutant siblings (Fig. 5F,F’). In two compound mutants, the walls of the ventricles remained highly trabeculated and had not undergone compaction (Fig. 5G,G’). An identical spectrum of ventral body wall, OFT and ventricular septal defects is observed in *Bmp2^-/+^;Bmp4^-/+^* compound mutants (Goldman et al. 2009; Uchimura et al. 2009).

**Table 2A.** Progeny from *Bmp7^R-GFlag/+^* and *Bmp2^-/+^* intercrosses

| Age | Wildtype | *Bmp2*<sup>-/+</sup> | *Bmp7*<sup>R-GFlag/+</sup> | *Bmp2*<sup>-/+</sup>; *Bmp7*<sup>R-GFlag</sup> | Total |
| --- | --- | --- | --- | --- | --- |
| P21** | 12 (50%) | 7 (29%) | 5 (29%) | 0 (0%) | 24 |
| E15.5-16.5 | 8 (26%) | 8 (26%) | 10 (32%) | 5 (16%) | 31 |
| E14.5 | 18 (37%) | 11 (23%) | 10 (20%) | 10 (20%) | 49 |

Data are presented as number (percent). Asterisks indicate that the observed frequency is significantly different than the expected frequency by Chi-square analysis (\*\*P<0.01).

To assess whether BMP4/7 heterodimers are required for development, we intercrossed *Bmp7^R-GFlag/+^* and *Bmp4^-/+^* mice. Compound mutants were present at predicted Mendelian ratios at E9.5-11.5, but were not recovered at E12.5 or later (Table 2B). *Bmp4^-/+^;Bmp7^R-GFlag/+^* embryos appeared grossly normal at E9.5, and expression of the BMP target gene *Nkx2.5* was intact in the heart (Supplemental Fig. S4C-F’). However, by E10.5 all eight embryos that were examined were smaller than littermates (Fig. 5I-L) and showed defects in heart development (Fig. 5I-L’). Specifically, whereas the hearts of wild type and single mutant siblings had morphologically distinguishable atria, ventricles, and OFT (Fig. 5I’-K’), *Bmp4^-/+^;Bmp7^R-GFlag/+^* hearts had a single common atrium (CA), and a smaller, malformed right ventricle and OFT (Fig. 5L’). In addition, expression of the BMP target gene *Nkx2.5* was reduced in the hearts of all *Bmp4^-/+^;Bmp7^R-GFlag/+^* embryos (Fig. 5L’) relative to littermates (Fig. 5I’-K’).

**Table 2B.** Progeny from *Bmp7^R-GFlag/+^* and *Bmp4^-/+^* intercrosses

| Age | Wildtype | *Bmp4*<sup>-/+</sup> | *Bmp7*<sup>R-GFlag/+</sup> | *Bmp4*<sup>-/+</sup>; *Bmp7*<sup>R-GFlag</sup> | Total |
| --- | --- | --- | --- | --- | --- |
| E12.5-14.5* | 9 (38%) | 8 (33%) | 7 (29%) | 0 (0%) | 24 |
| E11.5 | 5 (24%) | 4 (19%) | 5 (24%) | 7 (33%) | 21 |
| E10.5 | 5 (16%) | 7 (23%) | 11 (35%) | 8 (26%) | 31 |
| E9.5 | 14 (34%) | 11 (27%) | 7 (17%) | 9 (22%) | 41 |

Data are presented as number (percent). Asterisks indicate that the observed frequency is significantly different than the expected frequency by Chi-square analysis (\*P<0.05).

To test whether other class II BMPs can compensate for loss of BMP7 in the heterodimer pool in *Bmp7^R-GFlag^* heterozygotes, we intercrossed *Bmp7^R-GFlag/+^* and *Bmp7^-/+^* mice. *Bmp7^R-GFlag/-^* mutants were present at predicted mendelian ratios at E9.5-10.5 but were not recovered at E12.5 or later (Table 2C). *Bmp7^R-GFlag/-^* embryos appeared grossly normal (Supplemental Fig. S4G-J), and expression of the BMP target gene *Nkx2.5*, was intact in the heart at E9.5 (Supplemental Fig. S4G’-J’). By E10.5, however, all four *Bmp7^R-GFlag/-^* embryos that were analyzed were smaller (Fig. 5P) and their hearts showed a common atrium (CA), small right ventricle (RV), small malformed OFT and reduced expression of *Nkx2.5* (Fig. 5P’) relative to siblings (Fig. 5M-O’). By contrast, compound mutants heterozygous for the control *Bmp7^Flag^* allele in combination with *Bmp2^/+^, Bmp4^-/+^* or *Bmp7^-/+^* were adult viable and showed no gross phenotypic defects (Supplemental Table S2A-C). Collectively, these findings suggest that heterodimers consisting of BMP7 together with BMP2 and/or BMP4 are essential for many early developmental processes including ventral body wall closure and formation of the heart. In addition, other class II BMPs cannot fully compensate for BMP7 in the heterodimer pool in the heart.

**Table 2C.** Progeny from *Bmp7^R-GFlag/+^* and *Bmp7^-/+^* intercrosses

| Age | Wildtype | *Bmp7*<sup>-/+</sup> | *Bmp7*<sup>R-GFlag/+</sup> | *Bmp7*<sup>-/+</sup>; *Bmp7*<sup>R-GFlag</sup> | Total |
| --- | --- | --- | --- | --- | --- |
| E12.5-14.5* | 7 (29%) | 9 (38%) | 8 (33%) | 0 (%) | 24 |
| E10.5 | 9 (37%) | 5 (21%) | 6 (25%) | 4 (17%) | 24 |
| E9.5 | 4 (17%) | 7 (31%) | 6 (26%) | 6 (26%) | 23 |

Data are presented as number (percent). Asterisks indicate that the observed frequency is significantly different than the expected frequency by Chi-square analysis (\*P<0.05).

### Biochemical analysis reveals the existence of BMP4/BMP7 heterodimers in early embryos

To obtain biochemical evidence for heterodimer formation, BMP7 was immunoprecipitated from E11.5 *Bmp7^R-GFlag/+^* or *Bmp7^Flag/+^* protein lysates using antibodies directed against the Flag tag in the mature domain. The ability of BMP4 to co-purify with BMP7 was assayed by probing immunoblots of immunoprecipitates with antibodies specific for the mature domain of BMP4. As shown in Figure 6, the cleaved BMP4 mature ligand, as well as the BMP4 precursor protein co-immunoprecipitated with Flag-tagged BMP7 in lysates from *Bmp7^R-GFlag/+^* or *Bmp7^Flag/+^* embryos, but not from wild type littermates.

**Figure 6.**
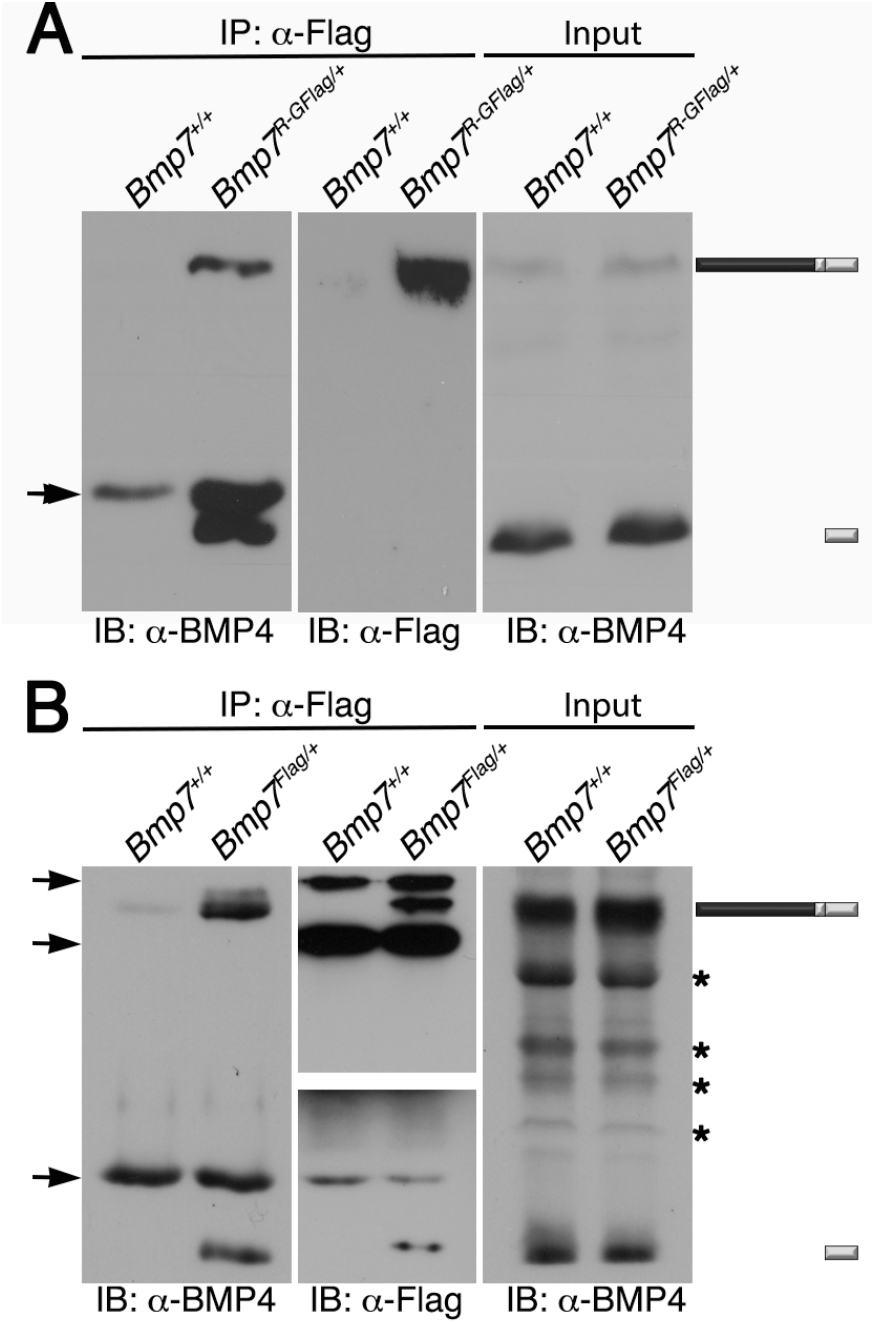
Endogenous BMP4 co-immunoprecipitates with BMP7. Antibodies specific for Flag-epitope tag were used to immunoprecipitate (IP) proteins from E11.5 *Bmp7*^+/+^, *Bmp7^R-GFlag/+^* or *Bmp7^Flag/+^* lysates. Immunoblots of IPs or total protein (input) were probed with antibodies specific for BMP4 or Flag as indicated below each panel. The position of precursor proteins and cleaved mature ligand is indicated on the right. Arrows indicates band corresponding to IgG heavy or light chains, asterisks mark non-specific bands. Results were reproduced in two independent experiments.

## Discussion

Previous studies have shown that heterodimers composed of BMP7 together with BMP2 or BMP4 have a higher specific activity than individual homodimers in specific in vitro assays, but it was unknown whether, or to what extent, endogenous class I/II BMP heterodimers are required for mammalian development. The current studies demonstrate that BMP2/7 and/or BMP4/7 heterodimers are the predominant functional signaling ligand in many tissues of early mouse embryos.

BMP activity is intact in the eye field of *Bmp7* null mutants at E9.5, but is absent in *Bmp7^R-GFlag^* homozygotes, suggesting that BMP7-containing heterodimers play an essential role in early inductive events in the eye. *Bmp4* and *Bmp7* are co-expressed in head surface ectoderm at the time of lens placode induction (E9), and both have been implicated in this process (Dudley and Robertson 1997; Furuta and Hogan 1998; Wawersik et al. 1999). Analysis of embryos engineered to express *Bmp4* or *Bmp6* from the *Bmp7* allele demonstrates that BMP6 can rescue eye defects in *Bmp7* null mutants, whereas BMP4 cannot (Oxburgh et al. 2005). This finding is consistent with the possibility that the higher specific activity of endogenous BMP4/7 heterodimers is essential to generate sufficient BMP activity for lens induction. In this scenario, another class II BMP (BMP6) could substitute for BMP7 to rescue heterodimer formation, whereas a class I BMP (BMP4) could not.

The role of BMP4/7 heterodimers in eye development is likely conserved in humans since *Bmp4* or *Bmp7* are co-expressed in the developing human eye, and mutations in either gene are associated with anophthalmia, microphthalmia and chorioretinal coloboma (Bakrania et al. 2008; Wyatt et al. 2010). Point mutations within the prodomain of BMP4 or BMP7 are also associated with an overlapping spectrum of brain and palate abnormalities (Bakrania et al. 2008; Suzuki et al. 2009; Wyatt et al. 2010; Reis et al. 2011), and several of these lead to single amino acid substitutions within short regions of the prodomain that are highly conserved between BMP4 and BMP7. We have shown that the prodomain of BMP4 is both necessary and sufficient to generate heterodimeric BMP4/7 ligands (Neugebauer et al. 2015), raising the possibility that the amino acid substitutions interfere with heterodimer formation.

The heart defects observed in *Bmp7^R-GFlag^* homozygotes appear earlier (by E9.5) but otherwise phenocopy those in *Bmp7^R-GFlag/+^;Bmp4^-/+^* and *Bmp7^R-GFlag/-^* mutants. This suggests that BMP4/7 heterodimers play essential roles in early stages of heart development that cannot be compensated for by other BMP family members. BMP4 is essential for septation of the ventricles, atrioventricular canal and outflow tract, as well as for valve formation and remodeling of the branchial arch arteries (Jiao et al. 2003; Liu et al. 2004; McCulley et al. 2008). Although BMP2 is not able to compensate for BMP4 during early heart development, our finding that *Bmp7^R-GFlag/+^;Bmp2^-/+^* compound heterozygotes have defects in the heart and in ventral body wall closure that phenocopy those observed in *Bmp2;Bmp4* compound heterozygotes (Goldman et al. 2009; Uchimura et al. 2009) suggest that BMP2 and BMP4 function redundantly as heterodimeric partners with BMP7 later in development. While *Bmp7* null mutants do not show defects in heart development, conditional deletion of both *Bmp7* and *Bmp4* from progenitors of the secondary heart field leads to persistent truncus arteriosus (Bai et al. 2013). Collectively, our results raise the possibility that these defects are due in part to reduction in the BMP4/7 and BMP2/7 heterodimer pool, rather than the loss of functionally redundant BMP4 and BMP7 homodimers.

Our observations that endogenous BMP activity is intact in the roof plate of the spinal cord in *Bmp7* null mutants at E9.5, but is reduced in *Bmp7^R-GFlag^* homozygotes suggest that heterodimers containing BMP7 are the physiologically relevant ligand(s) that are secreted from the surface ectoderm to induce the roof plate. *Bmp2, 4* and *7* are co-expressed in surface ectoderm overlying the neural tube by E8.5, whereas *Bmp5* and *Bmp6* are not expressed in this tissue at this stage (Dudley and Robertson 1997; Danesh et al. 2009). Furthermore, *Bmp5;Bmp6* and *Bmp5;Bmp7* double mutants show grossly normal dorsoventral patterning of the spinal cord (Solloway and Robertson 1999; Kim et al. 2001), indicating that signaling from the roof plate is intact.

One caveat to our conclusion that the phenotypic defects in *Bmp7^R-GFlag^* mutants are caused by loss of class I/II heterodimers is the possibility that BMP7R-GFlag forms inactive heterodimers with BMP5 and/or BMP6, effectively creating a double null mutant. While inactivation of BMP5 and/or BMP6 may indeed contribute to reduction in BMP activity in some tissues, including the heart, it cannot fully account for the defects observed in *Bmp7^R-GFlag^* homozygotes since they do not phenocopy those observed in *Bmp5;Bmp7* or *Bmp6;Bmp7* double mutants (Solloway and Robertson 1999; Kim et al. 2001).

The *Drosophila* BMP5-8 orthologs Screw and Glass bottom boat (GBB) undergo proteolytic processing at sites within the prodomain, in addition to cleavage of the site adjacent to the mature domain (Akiyama et al. 2012; Fritsch et al. 2012; Kunnapuu et al. 2014). In the case of GBB, cleavage of the prodomain site alone is sufficient to generate a bioactive ligand that signals at longer range than the conventional small ligand (Akiyama et al. 2012). Conserved PC consensus motifs can be identified within the prodomain of mammalian BMP7 (Akiyama et al. 2012), but the early lethality of *Bmp7^R-GFlag^* homozygotes demonstrates that cleavage at cryptic sites within the prodomain is not sufficient to generate functional BMP7 ligands that can support development. Furthermore, we are unable to detect BMP7 fragments generated by cleavage(s) at sites other than the previously identified PC motif in lysates from mouse embryos (current studies), *Xenopus* embryos or mammalian cells, or when BMP7 is cleaved by recombinant furin in vitro (Sopory et al. 2006; Neugebauer et al. 2015).

The endogenous BMP4 precursor is efficiently cleaved when dimerized with BMP7R-GFlag, raising questions as to how the mutant precursor blocks the function of wild type partners. BMP7 homodimers (Jones et al. 1994) and BMP4/7 heterodimers (Neugebauer et al. 2015) are secreted as a stable complex consisting of the cleaved mature ligand noncovalently associated with both propeptides. Type II BMP receptors compete with, and displace the BMP7 prodomain from the homodimer complex to initiate signaling (Sengle et al. 2008). It is possible that the uncleaved BMP7R-GFlag prodomain prevents the heterodimeric ligand from binding to or activating the receptor complex.

BMP4 and BMP7 preferentially form heterodimers rather than either homodimer when the two molecules are co-expressed in the same cell (Neugebauer et al. 2015). The current results suggest that this is a general theme for other class I and class II BMPs that are expressed in overlapping patterns. Because different BMPs are expressed at varying ratios in different cell types, the functional importance of heterodimers versus homodimers is likely to vary widely among different tissues and developmental stages. Additional biochemical and phenotypic analysis will be required to sort out which ligands are used in which tissues.

## Materials and Methods

### Mouse strains

Animal procedures followed protocols approved by the University of Utah Institutional Animal Care and Use Committees. *Bmp7^R-GFlag^* mice were generated using standard gene targeting procedures and *Bmp7^Flag^* mice were generated using CRISPR-Cas9 technology as described in detail in the Supplementary Materials and Methods. *Bmp4^LacZ/+^, Bmp2^-/+^* and BRE-LacZ mice were obtained from Dr. B. Hogan (Duke University), Dr. Y. Mishina (University of Michigan) and Dr. C. Mummery (Leiden University), respectively. *Bmp7^tm2Rob^* mice were obtained from Dr. E. Robertson (Cambridge University) and were used for all phenotypic analysis. *Bmp7^flox/flox^* mice were obtained from Dr. J. Martin (Baylor) and were crossed to CMV-cre mice to generate a null allele for analysis of BMP activity in BRE-LacZ crosses.

### Immunostaining, in situ hybridization and ß-galactosidase staining

For phosphoSmad immunostaining, E9.5 embryos were fixed in 4% paraformaldehyde in PBS at 4°C for one hour, incubated overnight in 30% sucrose in PBS at 4°C and then embedded in OCT (TissueTek). 10 μm cryosections were incubated overnight at 4°C with an antiphosphoSmad1/5/8 antibody (1:500; Cell Signaling 9511S) in PBS with 2% horse serum and 0.1% Triton X-100. Staining was visualized using anti-rabbit Alexa Fluor 488-conjugated secondary antibody (1:500; Molecular Probes). Embryos were processed for in situ hybridization with digoxigenin-labeled *Nkx2.5* riboprobes as described previously (Wilkinson and Nieto 1993). Quantitative in situ HCR was performed as described (Trivedi et al. 2018) using an *lmx1a* DNA probe set, a DNA HCR amplifier and hybridization, wash and amplification buffers purchased from Molecular Instruments. Whole mount mouse embryos were processed for in situ HCR as described (Choi et al. 2016). ß-galactosidase staining of *BRE-LacZ* embryos was performed as described (Lawson et al. 1999). Investigators were blinded to genotype until after morphology and/or staining intensity had been documented.

### Histology

Isolated embryos or dissected hearts were fixed in 4% paraformaldehyde in PBS, dehydrated and embedded in paraffin. Sections (10 μm) were stained with Hematoxylin and eosin.

### Transient transfection and Western blot analysis of cultured cells

HEK293T cells were plated on 10 cm culture dishes and transfected with 500 ng of DNA encoding BMP4, BMP7R-GFlag or empty vector. Cells were cultured for two days in serum containing media and then cultured for one additional day in serum free media before collecting conditioned media. HEK293T cells were incubated with conditioned- and control media for one hour, then cells were lysed and used for immunoblot analysis. Proteins were separated by electrophoresis on 10% or 12% SDS-polyacrylamide gels and transferred to PVDF membranes that were probed with anti-pSmad1/5/8 (Cell Signaling 9511S), anti-BMP4 (Santa Cruz sc12721), anti-HRP-conjugated Flag (Sigma F1804) or anti-ßactin (Abcam ab8229) primary antibodies followed by HRP-conjugated anti-rabbit IgG or HRP-conjugated anti-mouse IgG2b heavy chain specific (Jackson ImmunoResearch) secondary antibodies. Immunoreactive proteins were visualized using an ECL prime kit (GE HealthCare).

### Co-immunoprecipitation assays

Embryos were dissected from pregnant females at E11.5, homogenized in IP lysis buffer (150 mM NaCl, 20 mM Tris-Cl pH 7.5, 1mM EDTA, 1% Sodium deoxycholate, 1% NP40, 1X protease inhibitor (Thermo Scientific)) and protein concentration was measured using a BCA kit (Thermo Scientific). 1 mg of embryo lysate was diluted to 1 ml with IP lysis buffer pre-cleared by incubating with 100 μl protein A/G agarose for 2 hours at 4°C. Samples were spun for 5 seconds in a microfuge and 950 μl of supernatant was transferred to a new tube and incubated with agarose beads-conjugated to anti-Flag antibody (1:100; Sigma) for two hours at 4°C, followed by three 10 minute washes in IP buffer. Samples were spun for 5 seconds in a microfuge, supernatant was discarded and proteins were recovered in 40 ul 2X Laemmli sample buffer by boiling for 5 minutes prior to SDS-PAGE and immunoblot analysis.

### Skeletal preparations

Skeletal staining was performed as described (Hogan 1994).

## Acknowledgements

We thank Anne E. Martin for generating the schematic illustrations in Fig. 1 and Fig. 5, and Chris Gregg, Suzi Mansour and Rich Dorsky for helpful comments on the manuscript. This work was supported by the National Institutes of Health (RO1HD037976 to JLC, T32DK007115 to JMN, T32DK007115 to YSG, and T32HD007491 to AMN). This work utilized DNA, peptide, transgenic mouse and imaging shared resources supported by the Huntsman Cancer foundation and the National Cancer Institute of the NIH (P30CA042014) and the mutation generation and detection core supported in part by a grant from the National Institute of Diabetes and Digestive and Kidney Diseases (U54DK110858). The content is solely the responsibility of the authors and does not represent the official views of the NIH.

## Author contributions

H-S.K. and J.N. designed and performed experiments and interpreted data. H-S.K. and J.L.C. prepared figures and wrote the manuscript. A.M. maintained the mouse colony, genotyped mice and performed Southern blot analysis of ES cells. A.T. assisted with generating targeting constructs and performed Southern blot analysis of ES cells.

---

**Fig. S1.**
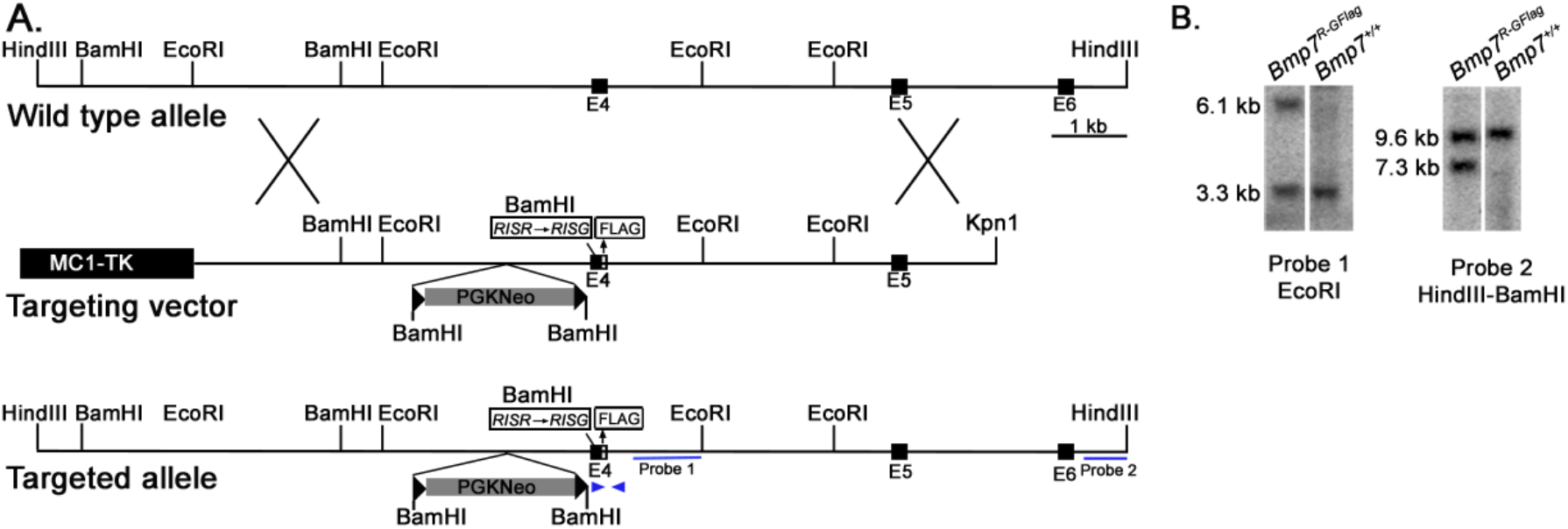
Generation of *Bmp7^R-GFlag^* mice. (A) Genomic organization of the wild type *Bmp7* allele, the targeting vector and the *Bmp7^R-GFlagNeo^* allele. The positions of the external (probe 2) and internal (probe 1) probes used for Southern analysis, and primers (arrows) surrounding the Flag tag that were used for PCR based genotyping are indicated. (B) Southern blot analysis of genomic DNA from targeted or non-targeted *(Bmp7^+/+^)* ES cells. Genomic DNA was digested with EcoRI or HindII and BamHI and hybridized with Probe 1 or Probe 2.

**Fig. S2.**
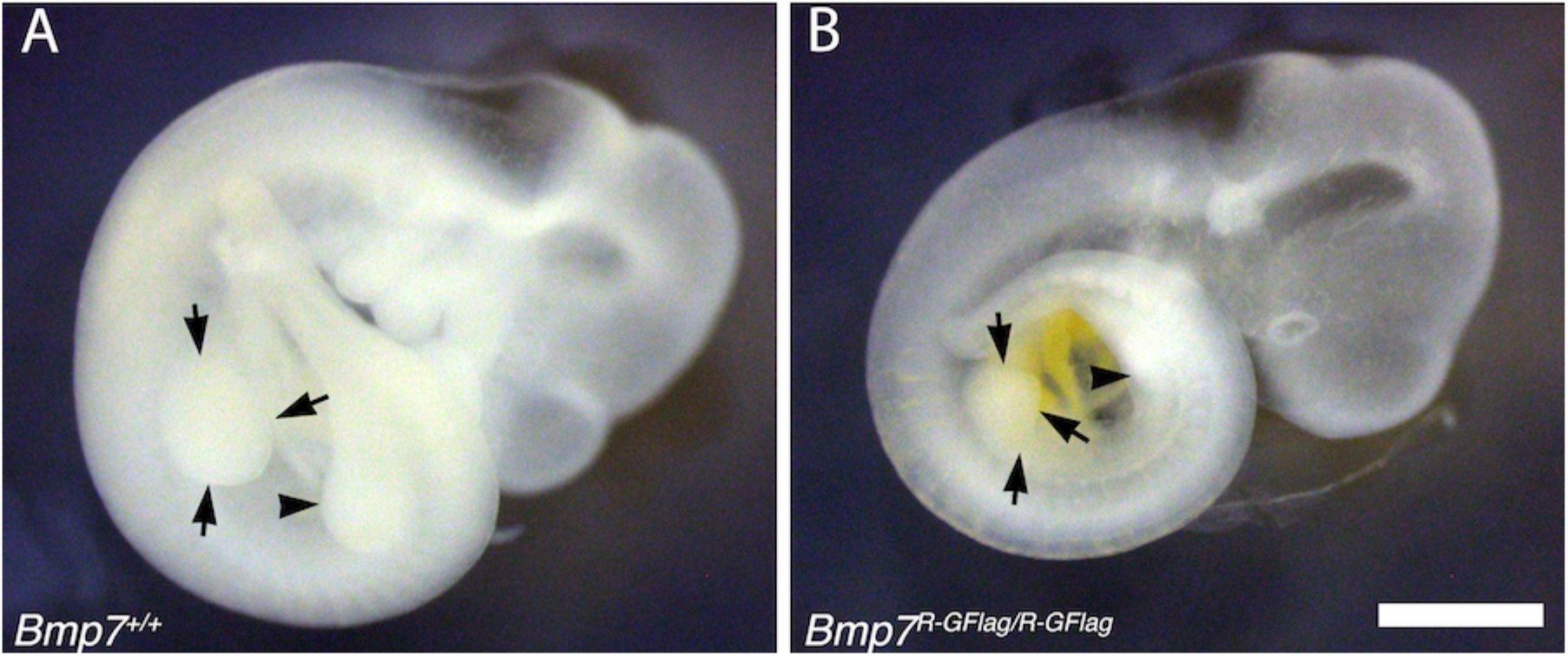
Limb bud size is reduced in *Bmp7^R-GFlag^* homozygotes. E9.5 wild type (A) and *Bmp7^R-GFlag/R-GFlag^* (B) littermates are shown. Arrows denote the forelimb bud and arrowhead denotes the hindlimb bud. Scale bar corresponds to 1 mm.

**Fig. S3.**
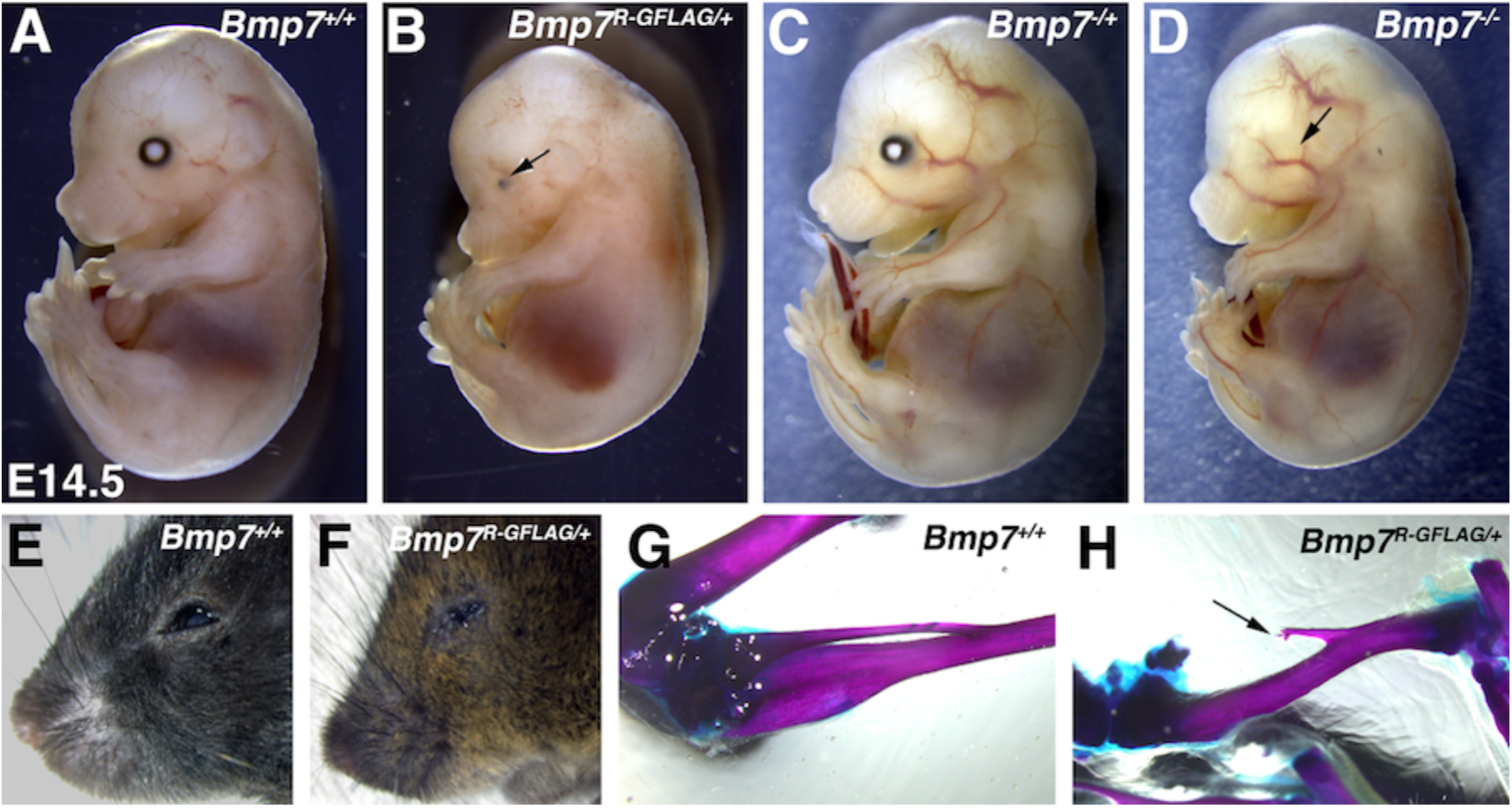
*Bmp7^R-GFlag^* heterozygotes show skeletal and eye defects that are absent in *Bmp7* null heterozygotes. (A-F) Photograph of E14.5 (A-D) and adult (E-F) wild type or *Bmp7* mutant embryos. Arrows denote small or absent eye in 3 out of 13 *Bmp7^R-GFlag/+^*, zero out of eight *Bmp7*^-/+^ and five out of five *Bmp7*^−/−^ mice analyzed between E14.5-18.5. (G-H) Representative skeletal preparations showing that the fibula (arrow) is shortened and not attached to the knee in two out of fourteen adult (3-5 months old) *Bmp7^R-GFlag^* heterozygotes that were examined.

**Fig. S4.**
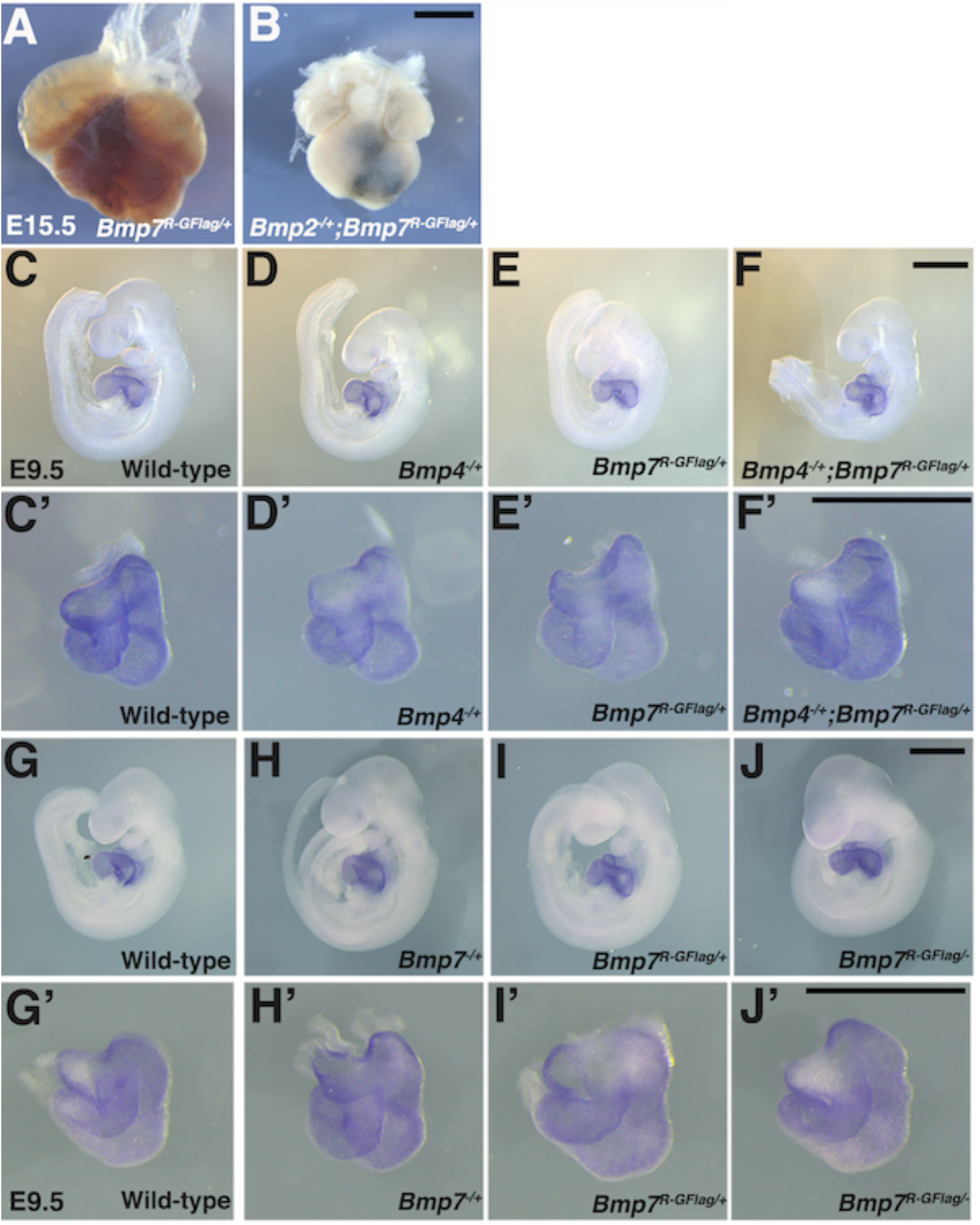
Heart defects are present in *Bmp2^-/+^;Bmp7^R-GFlag/+^* embryos at E15.5 and are absent in *Bmp4^-/+^;Bmp7^R-GFlag/+^* and *Bmp7^R-GFlag/-^* embryos at E9.5. (A-B) Photograph of heart dissected from *Bmp2^/+^;Bmp7^R-GFlag/+^* embryo and *Bmp7^R-GFlag/+^* littermate. (C-J’) Expression of *Nkx2.5* was analyzed by whole mount in situ hybridization in littermates generated by intercrossing *Bmp4^-/+^* and *Bmp7^R-GFlag/+^* (C-F’) or *Bmp7^-/+^* and *Bmp7^R-GFlag/+^* mice (G-J’). Photographs of intact embryos at E9.5 (C-J) and photographs of hearts dissected from corresponding embryos (C’-J’) are shown. Scale bars in all panels correspond to 1 mm.

