## Supplemental methods, figures, tables for "BMP7 functions predominantly as a heterodimer with BMP2 or BMP4 during mammalian embryogenesis"

### Supplementary Materials and Methods

#### Generation and genotyping of mice

The targeting vector used to generate *Bmp7*$^{R\text{-}GFlagNeo}$ mice was constructed from BAC clone bMQ298P20 purchased from Source Bioscience. This targeting construct (illustrated in Fig. S1A) includes: (a) sequence encoding an in frame Flag epitope tag within the mature domain located 24 amino acids downstream of the cleavage site (-EALRMDYKDDDDKASVAG-; Flag epitope underlined), (b) two point mutations in exon 4 that introduce an arginine to glycine amino acid change at the S2 cleavage site (RISR-RISG) and a new BamHI site, and (d) a neomycin selectable marker flanked by loxP sites upstream of exon 4. Linearized vector was electroporated into R1 ES cells and homologous recombinants were selected with G418 and gancyclovir. Correctly targeted ES cell clones were identified by Southern analysis using probes derived from genomic sequences located both internal and external to the targeting vector. Positive clones were expanded and mutations and epitope tag sequences were verified by sequencing DNA fragments PCR-amplified from genomic DNA. Heterozygous ES cells were injected into C57BL/6J blastocysts, and the resulting chimeras were mated with C57BL/6J females to obtain *Bmp7*$^{R\text{-}GFlagNeo}$ heterozygotes. Two independent mouse lines for each strain were mated to Cre deleter mice (Schwenk et al. 1995) to remove the neomycin gene.

*Bmp7*$^{Flag}$ mice were generated using CRISPR-Cas9 mutagenesis as described (Qin et al. 2016). sgRNA RNA (5'-CTCGGACCTACCTGCCACAC-3') was synthesized by in vitro transcription of an oligo-based template and was injected into C57BL/6J zygotes together with a single stranded donor DNA repair template (5'-CGCAGCCAGAATCGCTCCAAGACGCCAAAGAACCAAGAGGCCCTGAGGATGGACTACAAAGACGATGACGATAAAGCtAGcGTGGCAGgtaggtccgagcagctggaggggaccagctcattgcagatgctt-3'; sequence encoding FLAG epitope underlined) and Cas9 protein. G0 founders were crossed to

C57BL/6J females to obtain heterozygotes. DNA fragments PCR-amplified from genomic DNA were sequenced to verify the presence of the epitope tag and absence of other sequence changes. Genotypes were determined by PCR amplification of tail DNA using primers (illustrated in Fig. S1A) that anneal to sequence immediately surrounding the Flag epitope tag (5' primer: 5'-CAAGTTGGCAGGCCTGAT-3' and 3' primer: 5'- AAAGACACGTCCCAGGTCAC-3') under the following conditions: 94°C for 30 seconds, 60°C for 30 seconds, 72°C for 30 seconds, 35 cycles.

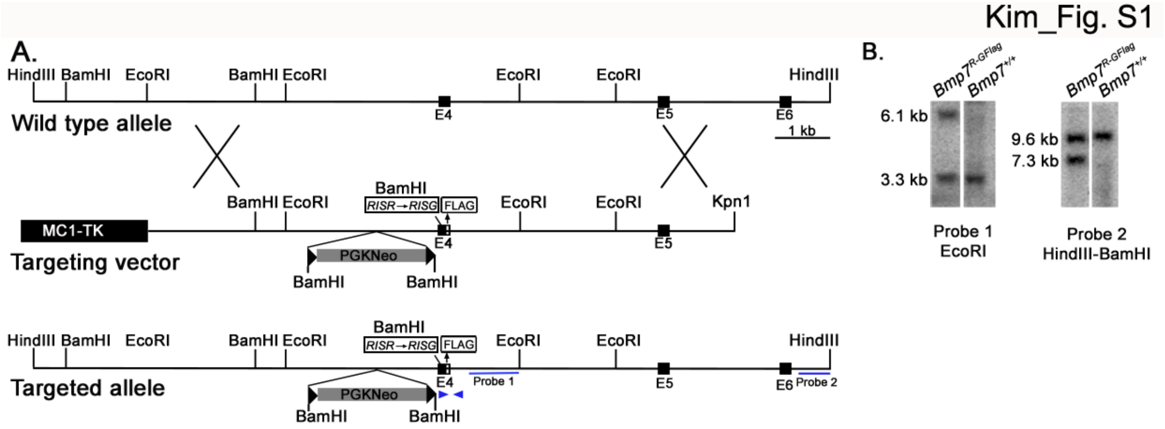


**Fig. S1. Generation of *Bmp7*$^{R\text{-}GFlag}$ mice.** (A) Genomic organization of the wild type *Bmp7* allele, the targeting vector and the *Bmp7*$^{R\text{-}GFlagNeo}$ allele. The positions of the external (probe 2) and internal (probe 1) probes used for Southern analysis, and primers (arrows) surrounding the Flag tag that were used for PCR based genotyping are indicated. (B) Southern blot analysis of genomic DNA from targeted or non-targeted (*Bmp7*$^{+/+}$) ES cells. Genomic DNA was digested with EcoRI or HindII and BamHI and hybridized with Probe 1 or Probe 2.

Kim_Fig. S2

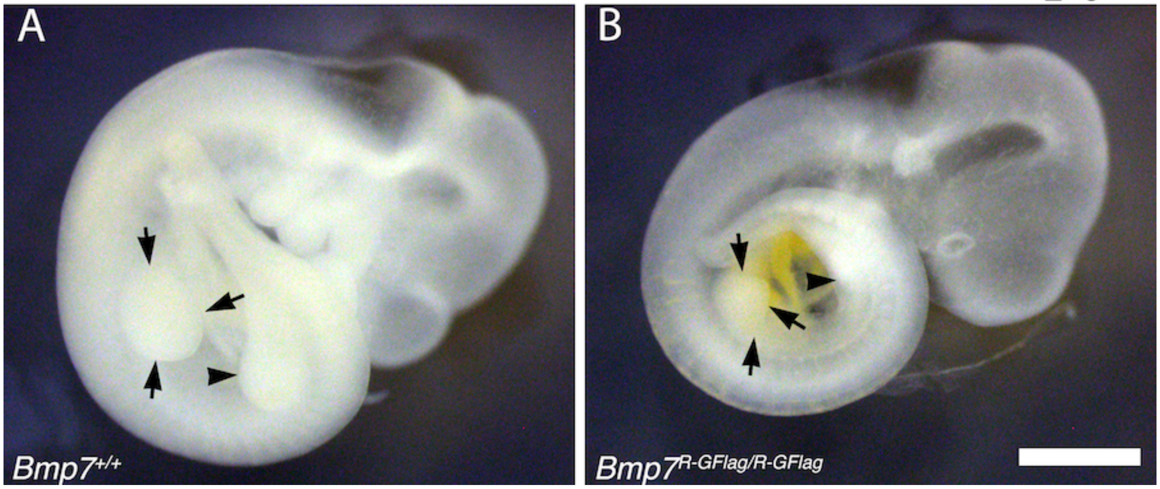


Fig. S2. Limb bud size is reduced in *Bmp7*$^{R\text{-}GFlag}$ homozygotes. E9.5 wild type (A) and *Bmp7*$^{R\text{-}GFlag/R\text{-}GFlag}$ (B) littermates are shown. Arrows denote the forelimb bud and arrowhead denotes the hindlimb bud. Scale bar corresponds to 1 mm.

Kim_Fig. S3

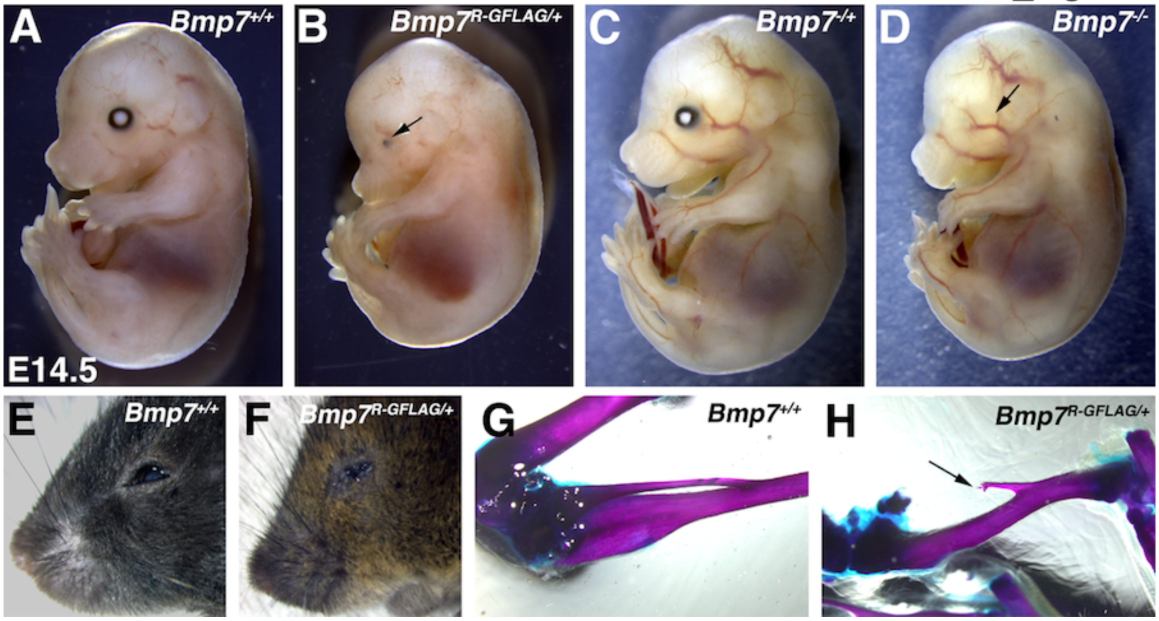


Fig. S3. *Bmp7*$^{R\text{-}GFlag}$ heterozygotes show skeletal and eye defects that are absent in *Bmp7* null heterozygotes. (A-F) Photograph of E14.5 (A-D) and adult (E-F) wild type or *Bmp7* mutant embryos. Arrows denote small or absent eye in 3 out of 13 *Bmp7*$^{R\text{-}GFlag/+}$, zero out of eight *Bmp7*$^{-/+}$ and five out of five Bmp7$^{-/-}$ mice analyzed between E14.5-18.5. (G-H) Representative skeletal preparations showing that the fibula (arrow) is shortened and not attached to the knee in two out of fourteen adult (3-5 months old) *Bmp7*$^{R\text{-}GFlag}$ heterozygotes that were examined.

Kim_Fig. S4

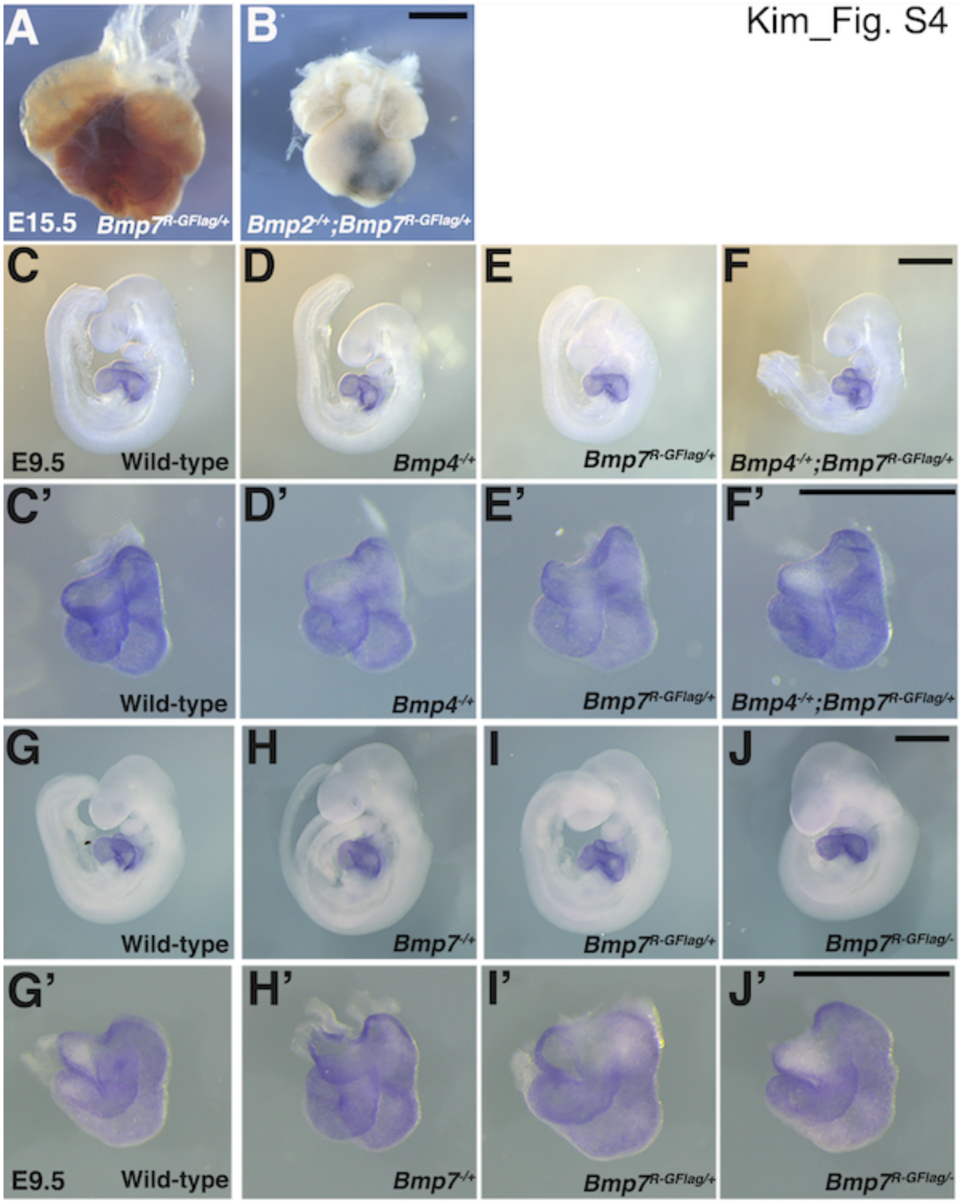


Fig. S4. Heart defects are present in *Bmp2*$^{-/+}$;*Bmp7*$^{R\text{-}GFlag/+}$ embryos at E15.5 and are absent in *Bmp4*$^{-/+}$;*Bmp7*$^{R\text{-}GFlag/+}$ and *Bmp7*$^{R\text{-}GFlag/-}$ embryos at E9.5. (A-B) Photograph of heart dissected from *Bmp2*$^{-/+}$;*Bmp7*$^{R\text{-}GFlag/+}$ embryo and *Bmp7*$^{R\text{-}GFlag/+}$ littermate. (C-J') Expression of *Nkx2.5* was analyzed by whole mount in situ hybridization in littermates generated by intercrossing *Bmp4*$^{-/+}$ and *Bmp7*$^{R\text{-}GFlag/+}$ (C-F') or *Bmp7*$^{-/+}$ and *Bmp7*$^{R\text{-}GFlag/+}$ mice (G-J'). Photographs of intact embryos at E9.5 (C-J) and photographs of hearts dissected from corresponding embryos (C'-J') are shown. Scale bars in all panels correspond to 1 mm.

**Table S1. Progeny from *Bmp7*$^{R\text{-}GFlag/+}$ and *Bmp7*$^{+/+}$ intercrosses**

| Age | *Bmp7*$^{+/+}$ | *Bmp7*$^{R\text{-}GFlag/+}$ | Total |
|---|---|---|---|
| **P28** | 155 (51%) | 151 (49%) | 306 |

Data are presented as number (percent).

**Table S2A. Progeny from *Bmp7*$^{Flag/+}$ and *Bmp2*$^{-/+}$ intercrosses**

| Age | Wildtype | *Bmp2*$^{-/+}$ | *Bmp7*$^{Flag/+}$ | *Bmp2*$^{-/+}$;*Bmp7*$^{Flag}$ | Total |
|---|---|---|---|---|---|
| P28 | 5 (21%) | 6 (25%) | 33 (29%) | 5 (21%) | 24 |

Data are presented as number (percent).

**Table S2B. Progeny from *Bmp7*$^{Flag/+}$ and *Bmp4*$^{-/+}$ intercrosses**

| Age | Wildtype | *Bmp4*$^{-/+}$ | *Bmp7*$^{Flag/+}$ | *Bmp4*$^{-/+}$;*Bmp7*$^{Flag}$ | Total |
|---|---|---|---|---|---|
| P28 | 4 (22%) | 4 (22%) | 5 (28%) | 5 (28%) | 18 |

Data are presented as number (percent).

**Table S2C. Progeny from *Bmp7*$^{Flag/+}$ and *Bmp7*$^{-/+}$ intercrosses**

| Age | Wildtype | *Bmp7*$^{-/+}$ | *Bmp7*$^{Flag/+}$ | *Bmp7*$^{-/+}$;*Bmp7*$^{Flag}$ | Total |
|---|---|---|---|---|---|
| P28 | 5 (29%) | 4 (23%) | 3 (18%) | 5 (29%) | 17 |

Data are presented as number (percent).
